# Mosaic RBD nanoparticles protect against multiple sarbecovirus challenges in animal models

**DOI:** 10.1101/2022.03.25.485875

**Authors:** Alexander A. Cohen, Neeltje van Doremalen, Allison J. Greaney, Hanne Andersen, Ankur Sharma, Tyler N. Starr, Jennifer R. Keeffe, Chengcheng Fan, Jonathan E. Schulz, Priyanthi N.P. Gnanapragasam, Leesa M. Kakutani, Anthony P West, Greg Saturday, Yu E. Lee, Han Gao, Claudia A. Jette, Mark G. Lewis, Tiong K. Tan, Alain R. Townsend, Jesse D. Bloom, Vincent J. Munster, Pamela J. Bjorkman

## Abstract

To combat future SARS-CoV-2 variants and spillovers of SARS-like betacoronaviruses (sarbecoviruses) threatening global health, we designed mosaic nanoparticles presenting randomly-arranged sarbecovirus spike receptor-binding domains (RBDs) to elicit antibodies against conserved/relatively-occluded, rather than variable/immunodominant/exposed, epitopes. We compared immune responses elicited by mosaic-8 (SARS-CoV-2 and seven animal sarbecoviruses) and homotypic (only SARS-CoV-2) RBD-nanoparticles in mice and macaques, observing stronger responses elicited by mosaic-8 to mismatched (not on nanoparticles) strains including SARS-CoV and animal sarbecoviruses. Mosaic-8 immunization showed equivalent neutralization of SARS-CoV-2 variants including Omicron and protected from SARS-CoV-2 and SARS-CoV challenges, whereas homotypic SARS-CoV-2 immunization protected only from SARS-CoV-2 challenge. Epitope mapping demonstrated increased targeting of conserved epitopes after mosaic-8 immunization. Together, these results suggest mosaic-8 RBD-nanoparticles could protect against SARS-CoV-2 variants and future sarbecovirus spillovers.

---

Two animal coronaviruses from the sarbecovirus lineage, severe acute respiratory syndrome coronavirus (SARS-CoV) and SARS-CoV-2 (hereafter SARS-1 and SARS-2), have caused epidemics or pandemics in humans in the past 20 years. SARS-2 triggered the COVID-19 pandemic that has been ongoing for over two years despite rapid development of effective vaccines (*1*). Unfortunately, new SARS-2 variants of concern (VOCs), including the heavily mutated Omicron VOCs (*2–7*), may prolong the COVID-19 pandemic. In addition, the discovery of diverse sarbecoviruses in bats, some of which bind the SARS-1 and SARS-2 entry receptor, angiotensin-converting enzyme 2 (ACE2) (*8–14*), raises the possibility of another coronavirus pandemic. Hence there is an urgent need to develop vaccines and therapeutics to protect against both SARS-2 VOCs and zoonotic sarbecoviruses.

Currently approved SARS-2 vaccines include the viral spike (S) trimer (*1*), consistent with S being the primary target of neutralizing antibodies (*15–24*). A coronavirus S trimer mediates entry into a host cell after one or more of its receptor-binding domains (RBDs) adopt an “up” position that allows interactions with a host cell receptor (Fig. 1A). Many of the most potent neutralizing antibodies against SARS-2 block binding ACE2 to the RBD (*16-20, 23-29*), and RBD targeting has been suggested for COVID-19 vaccine development (*30*). We classified neutralizing anti-RBD antibodies into four main classes (class 1, 2, 3, and 4) based on their epitopes and whether they recognized “up” and/or “down” RBDs on S trimers (*26*). Of note, the potent class 1 and class 2 anti-RBD antibodies, whose epitopes overlap with the ACE2 binding footprint, recognize a portion of the RBD that exhibits high sequence variability between sarbecoviruses (*26*). By contrast, the epitopes of class 4 antibodies, and to a somewhat lesser extent, class 3 antibodies, map to more conserved, but less accessible, regions of sarbecovirus RBDs (Fig. 1A). Substitutions in the RBDs of VOCs and variants of interest (VOIs) are also less common in the class 4 and class 3 regions (Fig. 1A), thus suggesting that a vaccine strategy designed to elicit class 3, class 4, and class 1/4 (class 4-targeting antibodies that block ACE2 binding (*31–33*)) could protect against potentially emerging zoonotic sarbecoviruses as well as current and future SARS-2 variants.

**Figure 1.**
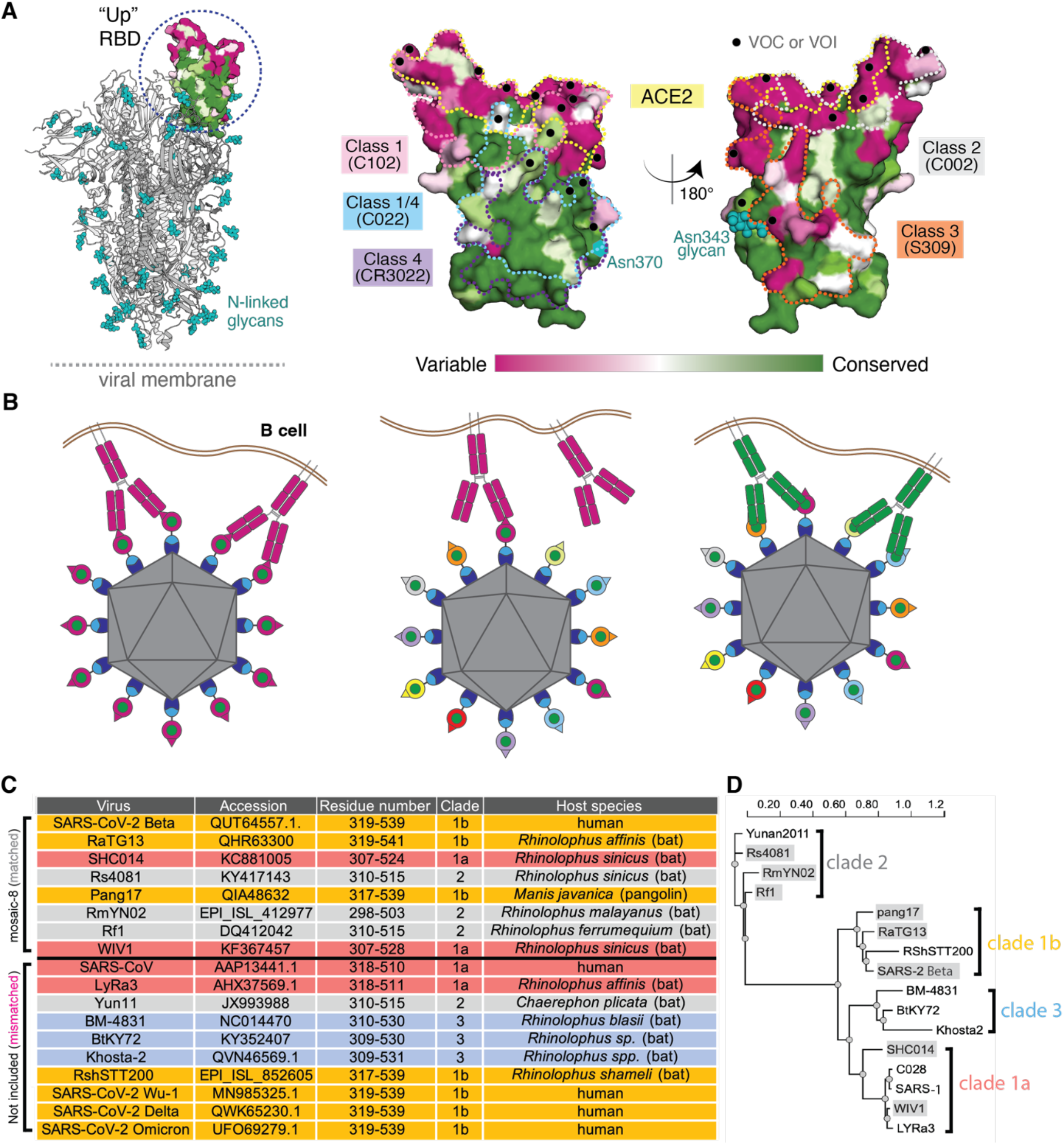
Mosaic nanoparticles may preferentially induce cross-reactive antibodies through avidity effects. (**A**) Left: Structure of SARS-2 S trimer (PDB 6VYB) showing one “up” RBD (dashed circle). Right: Sequence conservation of the 16 sarbecovirus RBDs in panel D calculated by the ConSurf Database (*79*) shown on two views of an RBD surface (PDB 7BZ5). The ACE2 binding footprint (PDB 6M0J) is outlined by a yellow dotted line. Locations of residues that are substituted in SARS-2 variants of concern (VOCs) and variants of interest (VOIs) as of March 2022 (https://viralzone.expasy.org/9556) are indicated as black dots. Class 1, 2, 3, 4, and 1/4 epitopes are outlined in different colored dotted lines using information from structures of representative monoclonal antibodies bound to RBD or S trimer (C102: PDB 7K8M; C002: PDB 7K8T, S309: PDB 7JX3; CR3022: PDB 7LOP; C022: PDB 7RKU). The N-linked glycan attached to RBD residue 343 is indicated by teal spheres, and the potential N-linked glycosylation site at position 370 in RBDs derived from sarbecoviruses other than SARS-2 is indicated by a teal circle. (**B**) Schematic showing hypothesis for how mosaic RBD-nanoparticles could induce production of cross-reactive antibodies. Left: Clustered membrane-bound B cell receptors bind with avidity to a strain-specific epitope (dark pink triangle) on dark pink antigens attached to a homotypic particle. Middle: B-cell receptors cannot bind with avidity to strain-specific epitope (triangle) on dark pink antigen attached to a mosaic particle. Right: B-cell receptors can bind with avidity to common epitope (green circle) presented on different antigens attached to a mosaic particle, but not to strain-specific epitopes (triangles). (**C**) Sarbecoviruses from which the RBDs in mosaic-8b RBD-mi3 were derived (matched) and sarbecoviruses from which RBDs were not included in mosaic-8b (mismatched). Clades are defined as in (*13*). The Wuhan-Hu-1 SARS-2 RBD was used in mosaic-8gm instead of the SARS-2 Beta RBD. (**D**) Phylogenetic tree of selected sarbecoviruses calculated using PhyML 3.0 (*80*) based on amino acid sequences of RBDs aligned using Clustal Omega (*81*). Viruses with RBDs included in mosaic-8b are highlighted in gray rectangles.

Here, we describe animal immunogenicity and virus challenge studies to evaluate mosaic-8 RBD-nanoparticles, a potential pan-sarbecovirus vaccine in which RBDs from SARS-2 and seven animal sarbecoviruses (*14*) were covalently attached to a 60-mer protein nanoparticle (*34*). The probability of two adjacent RBDs being the same is low for mosaic-8 RBD nanoparticles, an arrangement chosen to favor interactions with B cells whose receptors can crosslink between adjacent RBDs to preferentially recognize conserved, but sterically occluded, class 3, class 4, and class 1/4 RBD epitopes (Fig. 1B). By contrast, homotypic SARS-2 RBD-mi3, a nanoparticle including 60 copies of a SARS-2 RBD (*34, 35*), is more likely to engage B cells with receptors recognizing sterically accessible, but less conserved, class 1 and class 2 RBD epitopes (Fig. 1B). Antisera from animals primed and boosted with mosaic-8 or homotypic SARS-2 both showed neutralizing activity against SARS-2, but cross-reactivity against sarbecoviruses was more extensive in mosaic-8 antisera. In addition, while both mosaic-8 and homotypic SARS-2-immunized animals were protected from SARS-2 challenge, only the mosaic-8-immunized animals were protected from SARS-1 challenge. Finally, epitope mapping by deep mutational scanning of polyclonal antibodies showed preferential binding to conserved RBD epitopes for mosaic-8 antisera, but binding to the more variable class 2 RBD epitope for homotypic antisera, consistent with the hypothesized mechanism for elicitation of different classes of anti-RBD antibodies by mosaic versus homotypic RBD nanoparticles (Fig. 1B). These results highlight the potential for a mosaic nanoparticle approach to elicit more broadly protective antibody responses than homotypic nanoparticle approaches. We conclude that mosaic-8 RBD-nanoparticles show promise as a candidate vaccine to protect from SARS-2, present and future variants, and from zoonotic sarbecoviruses that could spill over into humans.

## Results

We used the SpyCatcher-SpyTag system (*36, 37*) to covalently attach RBDs with C-terminal SpyTag003 sequences to a 60-mer nanoparticle (SpyCatcher003-mi3) (*38*), to make either mosaic-8b (each nanoparticle presenting the SARS-2 Beta RBD plus seven other sarbecovirus RBDs attached to the 60 sites) or homotypic (each nanoparticle presenting 60 copies of the SARS-2 Beta RBD) RBD-mi3 nanoparticles (fig. S1A). In addition to the SARS-2 Beta RBD, the other RBDs in mosaic-8b nanoparticles were chosen from clade 1a, 1b, and 2 sarbecoviruses (clades as defined in (*13*)) (Fig. 1C,D, fig. S1A). Immune responses against these strains in immunized animals were considered “matched” since each was represented by an RBD on mosaic-8b. Sarbecoviruses from clades 1, 2, and 3 and SARS-2 RBDs other than SARS-2 Beta that did not have RBDs represented on mosaic-8b nanoparticles were considered “mismatched” in immunological assays and challenge experiments. SARS-1 was chosen as a mismatched strain to allow challenge experiments and because it uses human ACE2 as its host receptor (*39*) and can therefore be evaluated in ACE2-dependent pseudotyped neutralization assays, although we note that SARS-1 is closely related to WIV1, a clade 1a bat sarbecoviruses represented on the nanoparticles. Two versions of mosaic-8 were used for experiments: mosaic-8b RBD-mi3 (SARS-2 Beta RBD and seven animal sarbecovirus RBDs) (Fig. 1C,D) and mosaic-8gm (mosaic-8 with a Wuhan-Hu-1 SARS-2 RBD plus the seven zoonotic RBDs (*34*) in which N-linked glycosylation site sequons at RBD position 484 were introduced in the clade 1a and 1b RBDs to occlude class 1 and 2 RBD epitopes (fig. S1A). Mosaic-8 and homotypic SARS-2 Beta nanoparticles were purified via SEC (fig. S1B) and validated by SDS-PAGE to show near 100% conjugation efficiency (fig. S1C). Dynamic light scattering (DLS) and negative stain EM demonstrated that conjugated nanoparticles were monodisperse and exhibited a defined diameter (fig. S1D,E), and interactions of human ACE2 and monoclonal antibodies with known epitopes exhibited expected binding profiles (fig. S2).

To compare the efficacies of mosaic and homotypic RBD-mi3 nanoparticle immunizations, we evaluated immune responses and protection from viral challenge in K18-human ACE2 (K18-hACE2) transgenic mice (*40*) (Fig. 2,3). K18-hACE2 mice express human ACE2 driven by a cytokeratin promotor in epithelia, including airway epithelia cells where SARS-2 infections often start and recapitulate severe COVID-19 upon infection with SARS-2 (*40–42*). Viral challenges of K18-hACE2 mice result in extensive weight loss, and death usually results from SARS-2 or SARS-1 infection (*41*). We chose this lethal challenge model to evaluate the highest levels of potential protection, which might then be used to extrapolate to the expected efficacy of a vaccine in humans.

**Figure 2.**
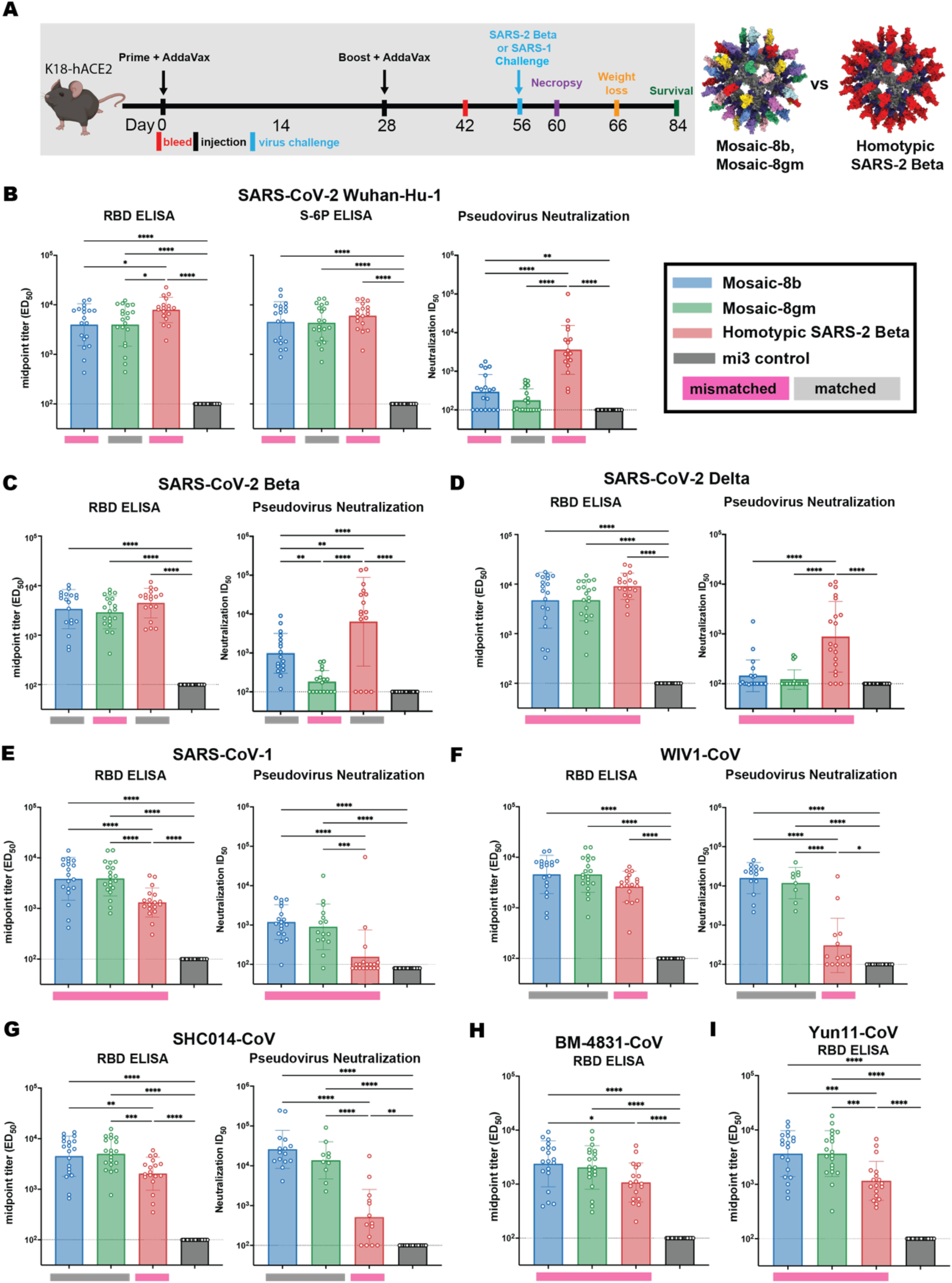
Mosaic-8b and homotypic SARS-2 Beta RBD-mi3 immunizations induced binding and neutralizing antibodies in K18 mice. (**A**) Left: Immunization schedule. K18-hACE2 mice were immunized with either mosaic-8b, mosaic-8gm, homotypic SARS-2 Beta, or unconjugated SpyCatcher003-mi3 nanoparticles. Right: Structural models of mosaic-8 and homotypic RBD-mi3 nanoparticles constructed using PDB 7SC1 (RBD), PDB 4MLI (SpyCatcher), and PDB 7B3Y (mi3). (**B-I**) ELISA and neutralization data from Day 42 (14 days post-Boost) for antisera from individual mice (open circles) presented as the mean (bars) and standard deviation (horizontal lines). ELISA results are shown as midpoint titers (EC_50_ values); neutralization results are shown as half-maximal inhibitory dilutions (ID_50_ values). Dashed horizontal lines correspond to the background values representing the limit of detection. Significant differences between cohorts linked by horizontal lines are indicated by asterisks: p<0.05 = *, p<0.01 = **, p<0.001 = ***, p<0.0001 = ****. Rectangles below ELISA and neutralization data indicate mismatched strains (pink; the RBD from that strain was not present on the nanoparticle) or matched strains (gray; the RBD was present on the nanoparticle).

**Figure 3.**
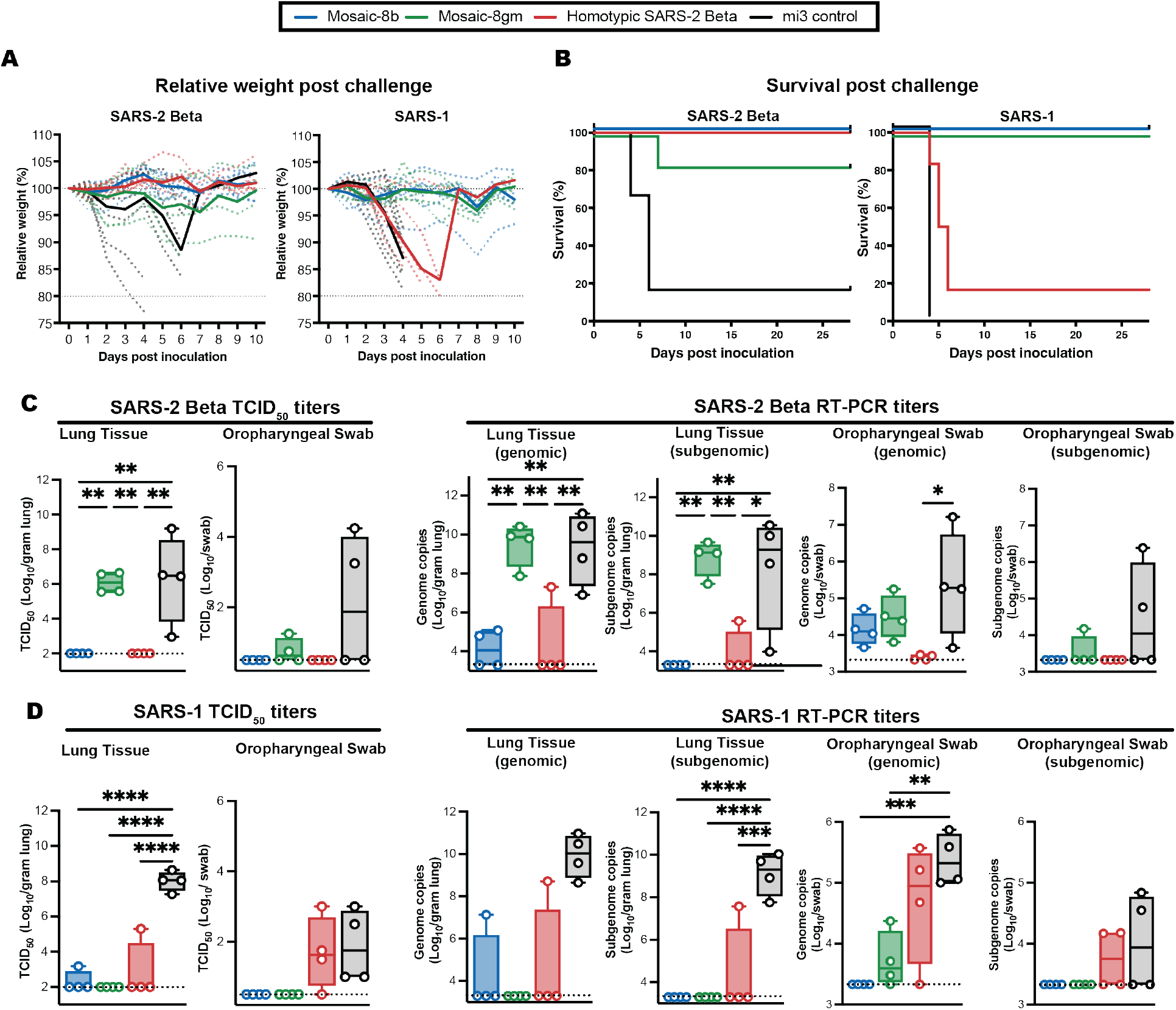
Mosaic-8b immunization protected against SARS-2 and SARS-1 challenges in K18-hACE2 mice, whereas homotypic SARS-2 immunization protected only against SARS-2. Mice were immunized and boosted with the indicated mi3 nanoparticles represented by different colors. (**A**) Weight changes after SARS-2 Beta or SARS-1 challenge. Mean weight in each vaccinated cohort indicated with a thick colored line. Weights of individual mice are indicated by colored dashed lines. (**B**) Survival after SARS-2 Beta or SARS-1 challenge. (**C**) Left: SARS-2 Beta infectious titers after challenge in lung tissue and oropharyngeal swabs. Right: Genomic and subgenomic SARS-2 Beta RNA copes determined by RT-PCR. (**D**) Left: SARS-1 infectious titers after challenge in lung tissue and oropharyngeal swabs. Right: Genomic and subgenomic SARS-1 RNA copies determined by RT-PCR. Significant differences between cohorts linked by horizontal lines are indicated by asterisks: p<0.05 = *, p<0.01 = **, p<0.001 = ***, p<0.0001 = ****.

K18-hACE2 mice were primed with either mosaic-8b, mosaic-8gm, homotypic SARS-2 Beta, or unconjugated SpyCatcher-mi3 nanoparticles adjuvanted with AddaVax and boosted 4 weeks later (Fig. 2A). In these experiments, SARS-2 Beta represented a matched sarbecovirus for the mosaic-8b RBD-mi3 and homotypic SARS-2 RBD-mi3 immunogens but was mismatched for mosaic-8gm. SARS-1 was mismatched for all three nanoparticle immunogens.

We first evaluated serum antibody responses in binding and pseudovirus neutralization assays 14 days after boosting (Fig. 2B-I). Serum ELISAs were conducted to assess binding to the indicated RBDs, and for Wuhan-Hu-1, also to the 6P stabilized version of soluble S trimer (*43*). ELISA titers against the Wuhan-Hu-1 RBD and S-6P were modestly higher for homotypic SARS-2 Beta immunized mice with respect to mosaic-8b and mosaic-8gm immunized mice, with all three immunogens eliciting high binding antibody titers (Fig. 2B). However, although the mosaic-8 immunogen antisera ELISA titers were not significantly different from homotypic RBD-mi3 antisera titers against the other SARS-2 variants (Fig. 2C,D), ELISA titers were increased against sarbecovirus RBDs derived from viruses other than SARS-2 (Fig. 2E-I); e.g., significantly higher binding titers when comparing the two mosaic-8 antisera versus the homotypic antisera against SARS-1 (mismatched; Fig. 2E), SHC014 (matched for the two mosaic-8 immunogens; Fig. 2G), BM48-31 (mismatched; Fig. 2H), and Yun11 (mismatched; Fig. 2I). As previously reported for RBD-mi3 immunogens (*34*), the trends for serum RBD binding were generally predictive of pseudovirus neutralization; however, the homotypic SARS-2 Beta antisera showed significantly higher neutralization titers against all three SARS-2 variants than the mosaic-8b and -8gm antisera (Fig. 2B-D). This contrasts with our previous reports of equivalent neutralization for antisera from mosaic-8 RBD-mi3 and homotypic SARS-2 RBD immunized mice (*34*). In those experiments, the SARS-2 RBD in both nanoparticles was derived from the Wuhan-Hu-1 strain rather than the Beta VOC, which could elicit increased levels of potent class 2 anti-RBD antibodies that bind RBDs with residue E484, which is substituted in most SARS-2 variants (*44*). Equivalent levels of anti-SARS-2 RBD binding antibodies, but lower neutralization potencies, in mosaic-8 versus homotypic RBD antisera are consistent with a larger portion of non-neutralizing antibodies against SARS-2 induced by mosaic-8 immunization. Non-neutralizing antibodies could be involved in protection since non-neutralizing antibodies have been shown to play a role in preventing severe COVID-19 from natural infection or challenges after vaccination (*45, 46*). Despite their reduced neutralization potencies against SARS-2, the mosaic-8b and -8gm antisera showed significantly higher neutralization titers than the homotypic antisera against clade 1a viruses such as SARS-1 (mismatched; Fig. 2E) and WIV1 and SHC014 (matched for mosaics; Fig. 2F,G).

The four groups of immunized K18-hACE2 mice (n=10) were challenged with SARS-2 Beta or SARS-1 (Fig. 2A). Four mice per group were euthanized at 4 days post challenge for viral load analysis, and the remaining six mice were monitored for survival up to 28 days post challenge. Mice in each cohort were evaluated for weight loss, survival, and levels of viral RNA (genomic and subgenomic) and infectious virus in lung tissue and oropharyngeal swabs (Fig. 3). Control animals immunized with unconjugated mi3 showed rapid weight loss and death four to six days after SARS-2 or SARS-1 challenge. As evaluated by relative weight loss, the mosaic-8b and homotypic SARS-2 Beta RBD-mi3 nanoparticles were equally protective against SARS-2 challenge, showing minimal to no weight loss, while some of the mosaic-8gm animals experienced transient weight loss but all but one recovered by 10 days post-challenge (Fig. 3A). By contrast, only the mosaic-8b and mosaic-8gm immunizations prevented weight loss after the SARS-1 challenge, while the homotypic SARS-CoV-2 Beta RBD-mi3 mice experienced similar weight loss as the mi3 control mice (Fig. 3A). The post challenge survival results were consistent with weight loss: after SARS-2 Beta challenge, both the mosaic-8b and homotypic immunized animals showed complete survival (100%), while five of six animals (∼83%) in the mosaic-8gm immunization group survived (Fig. 3B). After SARS-1 challenge, all but one of the mice in the homotypic SARS-2 Beta group reached endpoint criteria within six days (a delay compared to the control mi3 group, in which all animals reached endpoint criteria within four days), whereas all mice in the mosaic-8b and mosaic-8gm group survived the challenge during the 28 days of post-challenge monitoring (Fig. 3B). Altogether, despite elicitation of lower neutralization titers against SARS-2 compared to homotypic Beta RBD-mi3 antisera (Fig. 2B-D), immunization with mosaic-8b RBD-mi3 was fully protective against both matched (SARS-2 Beta) and mismatched (SARS-1) challenges in the K18-hACE2 mouse model. By contrast, immunization with homotypic SARS-2 RBD-mi3 was protective against the matched SARS-2 Beta challenge, but not against the mismatched SARS-1 challenge. Mosaic-8gm immunization protected against SARS-1 challenge but showed somewhat reduced efficacy against the SARS-2 Beta challenge, perhaps related to occluding the class 2 RBD epitope targeted by potent, but usually strain-specific, neutralizing antibodies.

We measured levels of infectious virus and viral RNA in lung and oropharyngeal swab samples from challenged K18-hACE2 mice (n=4), obtained at four days post challenge (Fig. 3C,D). Infectious virus titers were measured by determining the median tissue culture infectious dose (TCID_50_) of either oropharyngeal swab or lung tissue homogenate samples as described (*47*). In SARS-2 Beta-challenged mice, vaccination with either mosaic-8b or homotypic SARS-2 Beta completely suppressed viral replication in the lungs and oropharyngeal swabs, whereas levels of infectious virus in mosaic-8gm immunized animals were equivalent to control immunized mice in lungs. The SARS-2 Beta infectious viral load was lower for mosaic-8gm immunized mice with respect to control animals in the oropharyngeal swabs, suggesting partial protection in these animals (Fig. 3C; left). Almost all vaccinated animal groups displayed completely suppressed infectious SARS-1 in lungs compared to control immunized mice (Fig. 3D; left). However, only mosaic-8b and mosaic-8gm immunized animals showed complete suppression of infectious SARS-1 in oropharyngeal swabs, whereas homotypic SARS-2 Beta immunized animals showed infectious viral loads that were similar to control animals (Fig. 3D; left), possibly explaining the severity of SARS-1 in these animals (Fig. 3A,B).

Viral RNA copies in the lung tissue and oropharyngeal swabs were measured by RT-PCR using both genomic and subgenomic primer sets. Genomic RT-PCR titers reflect the overall RNA copies in the sample, including both replicating virus in infected cells and viral particles/debris, whereas subgenomic RNA is a better surrogate for infectious viral titers since it is produced in infected cells and poorly packaged into virions (*48*). In lung tissue samples, genomic and subgenomic SARS-2 Beta viral RNA titers were lower in mosaic-8b and homotypic SARS-2 Beta immunized animals compared to control immunized animals (Fig. 3C; right), consistent with infectious virus titer measurements (Fig. 3C; left panel) and protection against SARS-2 Beta challenge in mosaic-8b and homotypic SARS-2 Beta immunized animals (Fig. 3A,B). SARS-2 Beta RNA copies in oropharyngeal swabs were low for all immunized cohorts compared to the control (Fig.3C; right). Subgenomic SARS-1 viral RNA titers were completely suppressed in mosaic-8b and mosaic-8gm immunized animals in both lung tissue and oropharyngeal swabs with respect to the control (Fig. 3D; right), also consistent with infectious viral titers (Fig. 3C; left panel) and complete protection in these cohorts against SARS-1 challenge (Fig. 3A,B). The lack of suppression of viral RNA and infectious viral loads in oropharyngeal swabs from homotypic SARS-2 Beta immunized animals challenged with SARS-1 correlates with lethality from SARS-1 infection, possibly due to virus entry into the brain via nasal infection (*49*). Interestingly, homotypic SARS-2 immunized animals showed lower levels of subgenomic viral RNA (Fig. 3D; right, one animal in lung and 2 animals in oropharyngeal samples) with respect to control animals, suggesting partial control of SARS-1.

The K18-hACE2 mouse experiments were designed to evaluate survival, weight loss, and reduction or absence of viral replication as the primary metrics for vaccine efficacy (*49*). However, we also obtained lung tissue 4 days post challenge for analysis by hematoxylin and eosin (H&E) staining and immunohistochemistry (IHC) (Data S1). Upon challenge with SARS-2 Beta, two of four mi3 control animals exhibited lesions characterized by minimal interstitial pneumonia centered on terminal bronchioles and extending into the adjacent alveoli with minimal to moderate perivascular leucocyte cuffing and alveolar exudate (fig. S3B,D, Table S1, top). IHC for SARS-2 N protein showed staining in cells from three of four animals (<1-90% of type I and II cells) (fig. S3D,H, Table S1, top). In contrast, minimal to no lesions were observed in mosaic-8b and homotypic SARS-2 Beta immunized animals (fig. S3A,C, Table S1, top). Two of four animals immunized with mosaic-8gm exhibited lesions characterized by mild interstitial pneumonia. IHC staining for SARS-2 N protein showed minimal viral antigen present in mosaic-8b (1-5% of cells) and homotypic SARS-2 Beta immunized animals (two of four with <1% of stained cells) (fig. S3E,G, Table S1, top), whereas some staining was found in animals in the mosaic-8gm (three of four had <1-40% of stained type I and II pneumocytes). For the SARS-1 challenge, one animal in the control group exhibited lesions affecting 5% of the lung characterized by a minimal interstitial pneumonia centered on terminal bronchioles with minimal alveolar exudate (fig. S3L, Table S1, bottom), whereas all four animals exhibited antigen staining (10-90% staining of type I and II cells) (fig. S3P, Table S1, bottom). In contrast, animals immunized with mosaic-8b did not have any observable pulmonary pathology (fig. S3I, Table S1, bottom). One animal vaccinated with mosaic-8gm had minimal interstitial pneumonia with peribronchial inflammation affecting less than 1% of the lung (fig. S3J, Table S1, bottom). Animals vaccinated with homotypic SARS-2 Beta did not have pulmonary lesions, except for one animal with minimal perivascular cuffing (fig. S3K, Table S1, bottom). In addition, animals vaccinated with mosaic-8b, mosaic-8gm, and homotypic SARS-2 Beta showed minimal to no antigen staining (all <1% in type I and II pneumocytes) (fig. S3M-O, Table S1, bottom). Overall, mosaic-8b immunization was efficacious against both SARS-1 and SARS-2 challenge. Mosaic-8gm immunized animals showed low levels of viral antigen staining in lung tissue obtained from SARS-1 challenged animals, but not SARS-2 challenged animals, whereas homotypic SARS-2 Beta immunized animals showed control of SARS-2 Beta and SARS-1 in the lungs matching the suppression of viral load in lung tissue (Fig. 3C,D). Of note, viral control in lung tissue did not match severity of disease for vaccinated animals (Fig 3A,B), most likely due to the neurological basis of disease severity in this animal model (*49*).

To extend these results to another animal model of SARS-2 and SARS-1 infection, we also conducted immunization and challenge studies in non-human primates (NHPs). We evaluated only the most promising vaccine candidate, mosaic-8b RBD-mi3, comparing to a non-immunized cohort for challenges with either SARS-2 Delta or with SARS-1. Both challenge viruses were mismatched since the mosaic-8b nanoparticles did not include RBDs from SARS-2 Delta or SARS-1.

Eight NHPs were immunized and boosted at day 28 with mosaic-8b RBD-mi3 adjuvanted with VAC20 (2% aluminum hydroxide wet gel, Al_2_O_3_) (alum) and then boosted again at day 92 with mosaic-8b RBD-mi3 adjuvanted with MF59, a squalene-based oil in water emulsion adjuvant (*50*). Four weeks after the second boost, half of the NHPs in the vaccinated and control groups were challenged with SARS-2 Delta and the other half were challenged with SARS-1 (Fig. 4A).

**Figure 4.**
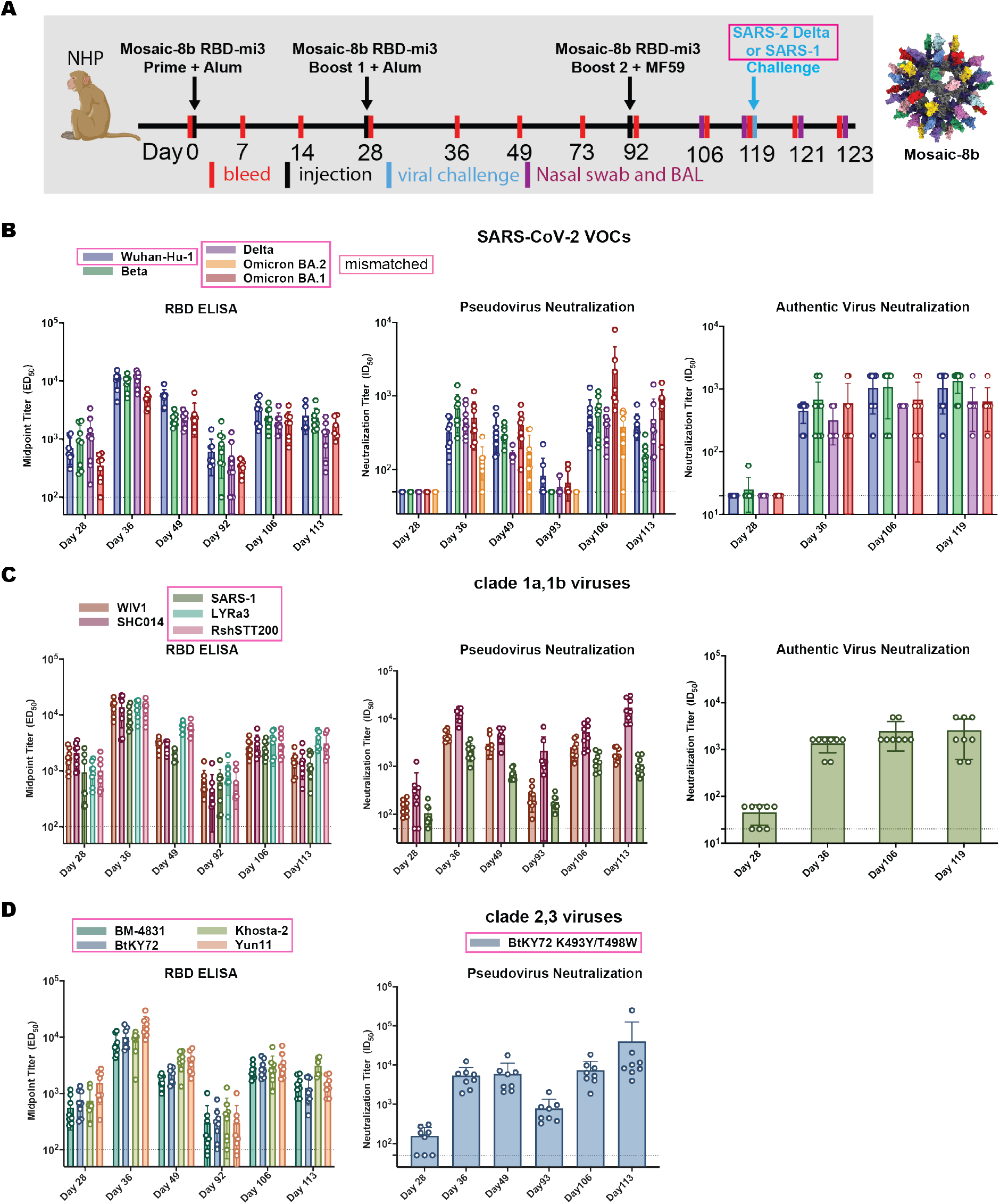
Mosaic-8b RBD-mi3 immunization induced binding and neutralizing antibodies in NHPs. Mismatched viruses are indicated by pink rectangular boxes. (**A**) Left: Immunization schedule. NHPs were primed and boosted with mosaic-8b RBD-mi3 in alum and boosted again with mosaic-8b RBD-mi3 in MF59. 8 immunized NHPs and 8 unimmunized NHPs were then challenged with either SARS-2 Delta (4 immunized and 4 unimmunized) or with SARS-1 (4 immunized and 4 unimmunized). Right: Structural model of mosaic-8b RBD-mi3 nanoparticles as shown in Fig. 2A. (**B-D**) Viruses for assays indicated as different colors; all were mismatched with respect to mosaic-8b RBD-mi3 except for SARS-2 Beta. ELISA and neutralization data for antisera from individual NHPs (open circles) presented as the mean (bars) and standard deviation (horizontal lines). ELISA results are shown as midpoint titers (EC_50_ values); neutralization results are shown as half-maximal inhibitory dilutions (ID_50_ values). Dashed horizontal lines correspond to the background values representing the limit of detection.

Polyclonal antisera were evaluated for binding to SARS-2 VOCs by ELISA and for neutralization activity using pseudovirus and authentic virus neutralization assays (Fig. 4B). The RBD ELISA and neutralization results showed similar trends, with relatively weak binding/neutralization before the first boost, rising levels after the first boost that contracted by Day 92, and then rising again and remaining above the 1:100 neutralization titers that correlate with ∼90% vaccine efficiency (*51*) after the second boost. Notably, the binding and neutralizing antibody levels were similar for SARS-2 Beta (matched) and the mismatched Wuhan-Hu-1, Delta, and Omicron BA.1 SARS-2 variants. We also evaluated binding and neutralization against RBDs and pseudoviruses from other sarbecovirus lineages (clade 1a, 2 and 3), including matched (WIV1, SHC014) and mismatched (SARS-1, LYRa3, RshSTT200, BM48-31, BtKY72, Khosta-2 and Yun11) viruses (Fig. 4C,D). Similar trends were observed for binding and neutralization of non-SARS-2 sarbecoviruses as seen for the SARS-2 variants: all RBDs were recognized by polyclonal antisera in ELISAs and neutralized in pseudovirus assays for human ACE2 entry-dependent viral strains for which neutralization assays could be conducted including mismatched strains: SARS-1 and a mutant form of BtKY72 (K493Y/T498W) that utilizes ACE-2 for entry (*13*). In all cases, the contracting antibody responses at Day 92 were restored by the second boost (Fig. 4C,D).

Mosaic-8b RBD-mi3 immunized and control NHPs were challenged with either SARS-2 Delta or SARS-1 (both mismatched) 28 days after the second boost (Fig. 4A). Protection was assessed by measuring infectious virus titers (Fig. 5A,B) and by viral RNA using RT-PCR (SARS-2 only; Fig. 5C) in bronchial alveolar lavage (BAL) and nasal swabs two or four days after challenge. We observed no detectable SARS-2 Delta infectious virus in BAL at either time point, whereas BAL from three of four control animals showed infectious SARS-2 virus (Fig. 5A). Nasal swabs showed low levels of virus in vaccinated animals, but at about two log significantly reduced levels compared to control NHPs (Fig. 5A), consistent with reports of detectable virus replication in upper airways in animals that were protected from clinical disease (*52*). For SARS-1 challenged animals, mosaic-8b immunized NHPs had no detectable viral titers in either BAL or nasal swabs, whereas all control animals had detectable viral titers (Fig. 5B; three of four in BAL and one of four in nasal swabs; individual animals identified by different colors). Although the low sample numbers precluded statistical significance of BAL or nasal swab differences in immunized versus control challenged animals, the lack of detectable SARS-1 virus in mosaic-8b immunized animals was suggestive of protection.

**Figure 5.**
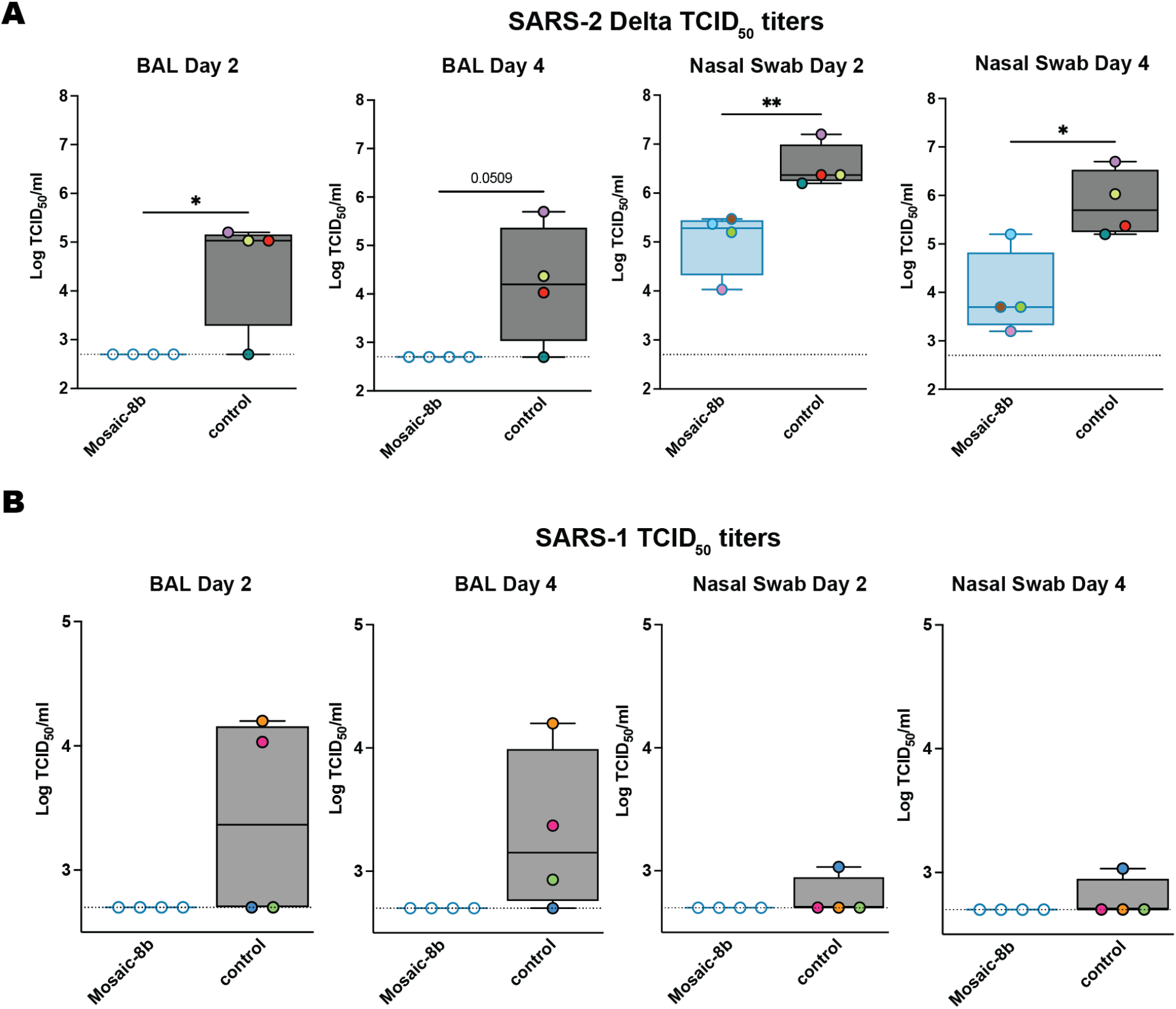
Mosaic-8b immunization protected NHPs against SARS-2 Delta and SARS-1 challenges. NHPs were immunized with mosaic-8b RBD-mi3 or not immunized (control) before challenge. (**A**) SARS-2 Delta infectious titers after challenge in BAL (left) and nasal swabs (right). Individual animals are denoted with different colors. (**B**) SARS-1 infectious titers after challenge in BAL (left) and nasal swabs (right). Individual NHPs in the unvaccinated control group are denoted with different colors to show that all four animals exhibited signs of detectable SARS-1 infectious virus in BAL and/or nasal swabs. Significant differences between cohorts linked by horizontal lines are indicated by asterisks: p<0.05 = *, p<0.01 = **, p<0.001 = ***, p<0.0001 = ****.

We next assessed whether mosaic and homotypic nanoparticles elicited different types of anti-RBD antibodies, as suggested by protection against matched challenge for both mosaic-8b and homotypic SARS-2 RBD-mi3 cohorts, but protection against mismatched challenge only for the mosaic-8b RBD-mi3 cohort. Immunizations with either mosaic-8b or homotypic SARS-2 Beta nanoparticles adjuvanted with AddaVax were conducted in BALB/c mice (prime and boost three weeks later) (fig. S4A). Anti-RBD antibodies in serum four weeks post boost evaluated by ELISA and neutralization (fig. S4B-G) exhibited similar characteristics as seen in the immunized K18-hACE2 mice (Fig. 2). To compare the characteristics of antibodies elicited by each type of RBD-nanoparticle, we mapped SARS-2 Beta epitopes recognized by immunization-elicited antibodies to investigate if mosaic-8b, but not homotypic SARS-2 Beta, preferentially elicited anti-RBD antibodies against conserved RBD epitopes as hypothesized (Fig. 1B). For these experiments, we used yeast-display deep mutational scanning to map mutations in the SARS-2 Beta RBD (*53, 54*) that escaped binding by antisera raised in BALB/c mice immunized with either mosaic-8b or homotypic SARS-2 Beta RBD-mi3 nanoparticles (Fig, 6; fig. S4,5).

The results showed that mosaic-8b antisera from six immunized mice primarily targeted more conserved RBD epitopes, including class 4 (RBD residues 383–386) and class 3 (residue 357) epitopes (Fig. 6A, fig. S6), whereas homotypic SARS-2 antisera primarily targeted variable RBD epitopes, particularly class 2 (residue K484) (Fig. 6B, fig. S6). These results confirmed that mosaic-8b RBD-mi3 elicited antibodies against the conserved class 3 and class 4 epitopes, as designed in the mosaic RBD nanoparticle approach (Figure 1A,B). By contrast, homotypic SARS-CoV-2 RBD-mi3 primarily elicited antibodies against the more variable class 2 epitope (characterized by RBD residue 484) that varies between sarbecoviruses and in SARS-2 VOCs (Fig. 1A).

**Figure 6.**
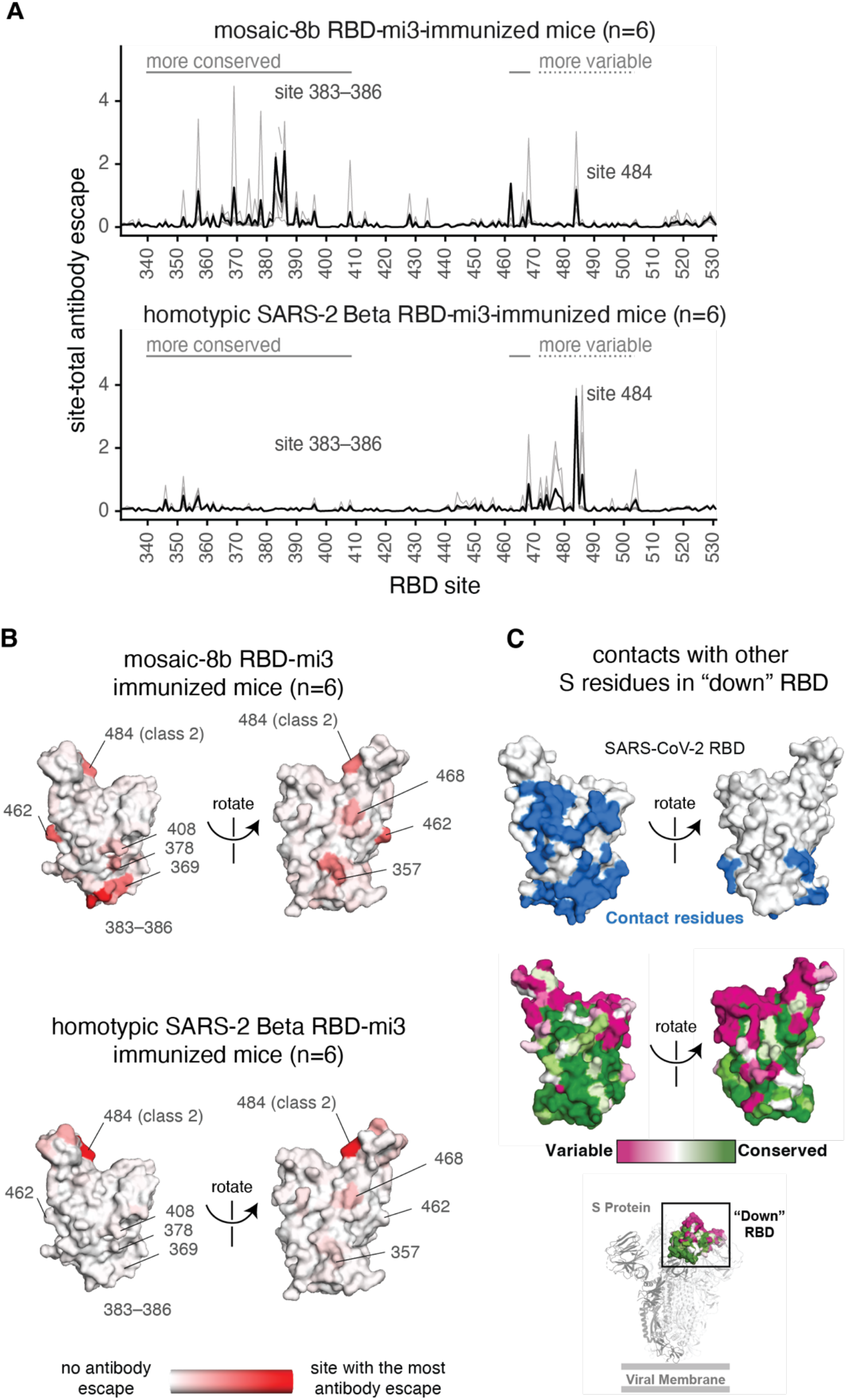
Antibodies elicited by mosaic-8b immunization map to conserved RBD epitopes, as compared to antibodies elicited by homotypic SARS-2 Beta immunization. (**A**) Deep mutational scanning was used to identify mutations that reduced binding of sera from BALB/c mice immunized with mosaic-8b RBD-mi3 (top) or homotypic SARS-2 Beta RBD-mi3 (bottom) to the SARS-2 Beta RBD. The y-axis shows the site-total antibody escape (sum of the antibody escape of all mutations at a site), with larger numbers indicating more antibody escape. Each light gray line represents one antiserum, and the heavy black lines indicate the average across the n=6 sera per group. RBD sites 340–408 and 462–468, which include the more conserved class 3/4 epitopes, are indicated with solid gray lines, and sites 472–503, which include sites from the more variable class 1/2 epitopes, are indicated with dashed lines. Note that the “conserved” and “variable” epitopes presented here were generalized for simple visualization and are not identical to more specific epitope-class definitions (*26, 59*). The highly variable RBD class 2 site 484 that is immunodominant among humans infected with SARS-2 (*44, 59*) and the subdominant class 4 sites 383–386 are labeled. (**B**) The average site-total antibody escape for mice immunized with mosaic-8b RBD-mi3 (top) or homotypic SARS-2 RBD-mi3 (bottom) mapped to the surface of the SARS-2 Beta RBD (PDB 7LYQ), with white indicating no escape, and red indicating sites with the most escape. Key sites are labeled, all of which are class 3/4 sites, except for the class 2 484 site. Interactive logo plots and structure-based visualizations of the antibody-escape maps are at https://jbloomlab.github.io/SARS-CoV-2-RBD_Beta_mosaic_np_vaccine/. Individual antibody-escape maps are in fig. S6; raw data are in Data S2 and at https://github.com/jbloomlab/SARS-CoV-2-RBD_Beta_mosaic_np_vaccine/blob/main/results/supp_data/all_raw_data.csv. (**C**) Top: Residues in a “down” RBD that contact other regions of spike shown in blue on an RBD surface (PDB 7BZ5). Interacting residues were identified using the PDBePISA software server (https://www.ebi.ac.uk/pdbe/prot_int/pistart.html) and the RBD from chain A of the spike trimer structure in PDB 7M6E. Middle: variable to conserved sarbecovirus sequence gradient (dark pink = variable; green = conserved) shown on RBD surface as in Fig. 1A. Bottom: Structure of SARS-2 S trimer (PDB 6VYB) showing “down” RBD (boxed) colored with the variable to conserved sarbecovirus sequence gradient.

We also mapped mutations that reduce binding of four serum samples from NHPs vaccinated with three doses of mosaic-8b RBD-mi3 (Day 106; Fig. 4). The NHP antibody-escape profiles were relatively broad, suggesting that no single mutation had a disproportionately large effect on binding (fig. S7). The NHP sera showed some skewing towards class 4 RBD epitopes and slight targeting of K484 (class 2) and T500 (class 3) (fig. S7). Differences in antibody-escape profiles for mosaic-8b immunized mice and NHPs could be related to species differences (*55*) and/or different immunization regimens (Fig. 2A, 4A). The broad escape profiles from NHP antisera may suggest either antibody binding to a broad set of RBD epitopes and/or a population of affinity-matured antibodies that are less affected by single point mutations. Nevertheless, the antibody-escape mapping results for mice and NHPs immunized with mosaic-8b RBD-mi3 are consistent with the hypothesis that the mosaic-8b nanoparticles elicit antibodies that target conserved RBD epitopes (Fig. 1B).

Mapping of sera from mosaic-8b-immunized mice and NHPs demonstrated relatively low, but non-zero, targeting of variable epitopes, e.g., typified by RBD residue 484. Of note, immunization with mosaic-8gm RBD-mi3 nanoparticles, in which class 1 and 2 epitopes in clade 1a and 1b RBDs were likely at least partially occluded by N-glycosylation at residue 484, was less protective against SARS-2 challenge than immunization with mosaic-8b, in which these epitopes were intact (Fig. 3). This result implies that retaining at least a subset of antibodies targeting the immunodominant class 1 and 2 epitopes may be important for protection against SARS-2 challenge. Strategies to occlude these epitopes by introducing N-glycans (*56, 57*) may thus impede optimal protection against SARS-2.

## Discussion

Anti-RBD antibodies raised by infection and vaccination can potently neutralize SARS-2 through blocking S trimer binding to the ACE2 receptor required for viral entry (*16-20, 23-29*). Although neutralizing antibodies recognize multiple RBD epitopes (*26, 58*), IgGs in human polyclonal plasmas tend to target the class 1 and class 2 epitopes that are undergoing rapid evolution in SARS-2 and that vary between zoonotic and human sarbecoviruses (*28, 44, 59*). Some monoclonal antibodies against these epitopes maintain breadth (*60*), but more commonly show partial or complete loss of potency against SARS-2 VOCs and only rarely cross-react with animal sarbecoviruses (*44, 61, 62*). By contrast, although less common and usually less potent than antibodies against class 1 and class 2 anti-RBD epitopes, antibodies against the more conserved class 3, 4, and 1/4 epitopes exhibit increased cross-reactivity across sarbecoviruses and SARS-2 VOCs (*27, 31-33, 63, 64*). Therefore, a vaccine that elicits such antibodies could serve to protect against SARS-2, its variants, and emerging zoonotic sarbecoviruses without the need for updating in the event of new VOCs and/or another sarbecovirus epidemic or pandemic.

Homotypic SARS-2 RBD or S trimer nanoparticles elicit potent neutralizing antibody responses that exhibit some degree of cross-reactivity across SARS-2 variants and sarbecoviruses (*34, 35, 65-74*). Here, we reproduce and extend those results in challenge studies demonstrating protection against SARS-2, including a mismatched variant, using a mosaic-8 RBD nanoparticle vaccine candidate. We demonstrated protection from SARS-2 challenge in animals immunized with homotypic SARS-2 RBD-mi3 and with mosaic-8b RBD-mi3 (challenges for the mosaic-8b including both a matched SARS-2 variant and a mismatched variant), despite mosaic-8b containing 1/8 as many SARS-2 RBDs as its homotypic SARS-2 counterpart. These results suggest that a mosaic RBD nanoparticle could be used now as a COVID-19 vaccine option to protect from current and future SARS-2 variants. Importantly, we also showed that mosaic-8b, but not homotypic SARS-2 RBD-mi3 nanoparticles, protected K18-hACE2 mice against lethality with a mismatched SARS-1 challenge, suggesting that a mosaic nanoparticle vaccine could also protect from disease caused by future mismatched, and heretofore unknown, zoonotic sarbecoviruses that could infect humans. It should be noted that, although homotypic SARS-2 RBD-mi3 nanoparticles did not confer complete protection against SARS-1, viral loads in lung tissue obtained from immunized animals were greatly reduced compared to control animals: only one of four animals had detectable viral loads in the lungs compared to four of four in the control group, demonstrating that some level of protection was achieved. Interestingly, a similar outcome was reported upon vaccination of aged mice with RBD-scNP, a homotypic SARS-2 RBD-conjugated ferritin nanoparticle that induced neutralizing antibodies against SARS-2 and pre-emergent sarbecoviruses: upon subsequent challenge with mouse-adapted SARS-1, viral loads were significantly reduced, but not absent, in lung tissue (four of five vaccinated animals exhibited reduced, but detectable SARS-1 virus, as compared with five of five with higher viral loads in the control group) (*74*).

The presence of immunodominant epitopes on viral antigens have contributed to preventing development of a universal influenza vaccine and vaccines against other antigenically-variable viruses such as HIV-1 and hepatitis C (*75*). Presentation of related, but antigenically different, viral antigens on a mosaic nanoparticle, a method to subvert immunodominance (*76*), is appropriate for making a pan-sarbecovirus vaccine because the SARS-2 S trimer contains immunodominant epitopes within its RBD that limit the breadth of antibodies elicited by SARS-2 infection or vaccination. Our demonstrations that a mosaic RBD nanoparticle presenting 8 different sarbecovirus RBDs protected against challenges from both matched and mismatched sarbecoviruses, as compared with homotypic SARS-2 RBD nanoparticles that protected fully only against a matched challenge, are consistent with RBD mapping experiments demonstrating that mosaic-8b, but not homotypic SARS-2 RBD-mi3 nanoparticles, primarily elicited antibodies against conserved RBD regions rather than the immunodominant class 1 and class 2 RBD epitopes. By including 8 different RBD antigens arranged randomly on a 60-mer nanoparticle, as compared to a smaller number of different RBDs that are not arranged randomly (*77*), the chances of stimulating production of cross-reactive antibodies against conserved regions is maximized because adjacent antigens are unlikely to be the same (*76*). The plug-and-display approach facilitated by SpyCatcher-SpyTag methodology (*36, 78*) allows straightforward production of mosaic nanoparticles with different RBDs attached randomly. Such nanoparticles could be used to protect against COVID-19 and future sarbecovirus spillovers and easily adapted to make other pan-coronavirus vaccines; for example, against MERS-like betacoronaviruses and/or against alpha or delta coronaviruses. Given the recent plethora of SARS-2 VOCs and VOIs that may be arising at least in part due to antibody pressure, a relevant concern is whether more conserved RBD epitopes might be subject to substitutions that would render vaccines and/or monoclonal antibodies targeting these regions ineffective. Although direct proof remains to be established, this scenario seems unlikely, as RBD regions conserved between sarbecoviruses and SARS-2 variants are generally involved in contacts with other regions of spike trimer (Fig. 6C) and therefore less likely to tolerate selection-induced substitutions.

## Supporting information

Pathology summary

## Acknowledgments

We thank Jost Vielmetter and the Caltech Protein Expression Center for assistance with protein production, Mark Howarth (Oxford) and Sumi Biswas (SpyBiotech) for helpful discussions about SpyCatcher-SpyTag methodology, Francis Laurent and Ruben Caputo (SPI Pharma) and Harshet Jain (Panacea Biotec) for VAC20 and MF59 adjuvants, respectively, Alexandra Walls and David Veesler (University of Washington) for a modified BtKY72 S gene for pseudovirus neutralization assays, Myndi Holbrook, Emmie de Wit, Brandi Williamson, Andrew Pekosz, Craig Martens, Kent Barbian, Stacey Ricklefs, Sarah Anzick, Ricki Feldmann, and the Public Health Agency of Canada for viruses used at RML, Kestrel Almquist, Allison Darrow, Kaitlyn Bauer, Amanda Weidow, Richard Cole, Lisa Heaney, Maarit Culbert, Brandon Bailes, Corey Henderson, Shane Gallogly, Lydia Crawford, and Taylor Lippincott for animal care at RML, and Claude Kwe Yinda (RML) for technical support. The following reagents were obtained through BEI Resources, NIAID, NIH: Cercopithecus aethiops Kidney Epithelial Cells Expressing Transmembrane Protease, Serine 2 and Human Angiotensin-Converting Enzyme 2 (Vero E6-TMPRSS2-T2A-ACE2), NR-54970. We thank Labcorp Drug Development–Antibody Reagents and Vaccines (Denver, PA) (formerly Covance, Inc.) for carrying out BALB/c mice immunizations for the RBD epitope mapping study.

## Funding

Wellcome Leap (PJB)

Bill and Melinda Gates Foundation INV-034638 (PJB); INV-004949 and INV-016575 (JDB)

Caltech Merkin Institute (PJB)

George Mason University Fast Grant (PJB)

Intramural Research Program of the National Institute of Allergy and Infectious Diseases (NIAID), National Institutes of Health (NIH) (1ZIAAI001179-01) (VJM)

NIAID/NIH contract numbers HHSN272201400006C and 75N93021C00015 (JDB)

National Institutes of Health grant R01AI141707 (JDB)

ORIP grant S10OD028685 (The Scientific Computing Infrastructure at Fred Hutch)

Caribbean Primate Research Center (Grants P40 OD012217 and 2U42OD021458 from ORIP/OD/NIH) (MGL)

TNS is a Howard Hughes Medical Institute Fellow of the Damon Runyon Cancer Research Foundation (DRG-2381-19).

JDB is an Investigator of the Howard Hughes Medical Institute.

TKT is funded by the Townsend-Jeantet Charitable Trust (charity number 1011770) and the EPA Cephalosporin Early Career Researcher Fellowship.

## Author contributions

Conceptualization: AAC, TKT, ART, PJB

Methodology: AAC, NvD, AJG, HA, AS, TNS, JRK, JES, MGL, JDB, VJM, PJB

Investigation: AAC, NvD, AJG, HA, AS, JRK, CF, JES, PNPG, LMK, APW, GS, YEL, HG, CAJ

Visualization: AAC, CF, AJG, JES, JDB Funding acquisition: PJB, JDB, VJM

Project administration: JRK, MGL, JDB, VJM, PJB

Supervision: AAC, NvD, HA, AS, JRK, APW, MGL, JDB, VJM, PJB

Writing – original draft: AAC, PJB

Writing – review & editing: AAC, NvD, AJG, HA, JRK, CF, TKT, ART, TNS, JDB, VJM, PJB

## Data and materials availability

All data are available in the main text or the supplementary materials. Raw Illumina sequencing data for the antibody-escape mapping experiments are available on the NCBI Short Read Archive (SRA) at BioProject PRJNA770094, BioSample SAMN26315988. Antibody-escape scores are available at https://github.com/jbloomlab/SARS-CoV-2-RBD_Beta_mosaic_np_vaccine/blob/main/results/supp_data/all_raw_data.csv. Materials are available upon request to the corresponding author with a signed material transfer agreement. This work is licensed under a Creative Commons Attribution 4.0 International (CC BY 4.0) license, which permits unrestricted use, distribution, and reproduction in any medium, provided the original work is properly cited. To view a copy of this license, visit https://creativecommons.org/licenses/by/4.0/. This license does not apply to figures/photos/artwork or other content included in the article that is credited to a third party; obtain authorization from the rights holder before using such material.

## MATERIALS AND METHODS

### Protein Expression

Mammalian expression vectors encoding the RBDs of SARS-2 Beta (GenBank QUT64557.1), SARS-CoV-2 Wuhan-Hu-1 (GenBank MN985325.1), RaTG13-CoV (GenBank QHR63300), SHC014-CoV (GenBank KC881005), Rs4081-CoV (GenBank KY417143), pangolin17-CoV (GenBank QIA48632), RmYN02-CoV (GSAID EPI_ISL_412977), Rf1-CoV (GenBank DQ412042), W1V1-CoV (GenBank KF367457), Yun11-CoV (GenBank JX993988), BM-4831-CoV (GenBank NC014470), BtkY72-CoV (GenBank KY352407), Khosta-2 CoV(QVN46569.1), RsSTT200-CoV (EPI_ISL_852605), LYRa3 (AHX37569.1) and SARS-CoV S (GenBank AAP13441.1) with an N-terminal human IL-2 or Mu phosphatase signal peptide were constructed as previously described (*34, 82*). 5 of the 8 RBD genes (SARS-2, RaTG13, pang17, WIV1, and SHC014) used to make mosaic-8gm were altered by site-directed mutagenesis to include a potential N-linked glycosylation site (PNGS) (N at position 484 and T at position 486). Each RBD was expressed to include a C-terminal hexahistidine tag (G-HHHHHH) and SpyTag003 (RGVPHIVMVDAYKRYK) (*38*) (for coupling to SpyCatcher003-mi3) or only a 15-residue Avi-tag (GLNDIFEAQKIEWHE) followed by a 6xHis tag (for ELISAs). RBDs were purified from transiently-transfected Expi293F cell (Gibco) supernatants by Ni-NTA and size-exclusion chromatography (SEC) as described (*82*), and RBDs with an introduced PNGS used for making mosaic-8gm RBD-mi3 were compared to their counterpart RBDs by SDS-PAGE to verify addition of extra N-glycans. SEC RBD fractions identified by SDS-PAGE were pooled and stored at 4°C or frozen in liquid nitrogen and stored at -80°C for longer term storage. A soluble SARS-2 trimer with 6P stabilizing mutations (*83*) was expressed and purified as described (*26*). Monoclonal human IgGs and human ACE-2 fused to human IgG Fc (hACE2-Fc) was expressed and purified as described (*26, 32*).

### Preparation of RBD-mi3 nanoparticles

SpyCatcher003-mi3 nanoparticles (*78*) were expressed in BL21 (DE3)-RIPL *E coli* (Agilent) transformed with the pET28a His6-SpyCatcher003-mi3 gene (Addgene) as described (*34, 84*). Briefly, transformed bacterial cell pellets were lysed in the presence of 2.0 mM PMSF (Sigma). Lysates were spun at 21,000xg for 30 min, filtered with a 0.2 µm filter, and mi3 particles were isolated by Ni-NTA chromatography using a HisTrap^TM^ HP column (GE Healthcare). Eluted particles were concentrated using an Amicon Ultra 15 mL 30K concentrator (MilliporeSigma) and SEC purified using a HiLoad® 16/600 Superdex® 200 (GE Healthcare) column equilibrated with 25 mM Tris-HCl pH 8.0, 150 mM NaCl, 0.02% NaN_3_ (TBS). SpyCatcher003-mi3 particles were stored at 4°C for up to 1 month and used for conjugations after 0.2 µm filtering or spinning at 21,000xg for 10 min.

Purified SpyCatcher003-mi3 nanoparticles were incubated with a 2-fold molar excess (RBD to mi3 subunit) of SpyTagged RBD (either a single RBD for homotypic SARS-2 RBD particles or an equimolar mixture of eight RBDs for mosaic particles) overnight at room temperature in TBS. The nanoparticles included the following RBDs: SARS-2 Beta (homotypic RBD-mi3); SARS-2 Beta, RaTG13, SHC014, Rs4081, RmYN02, pang17, Rf1, and WIV1 (mosaic-8b RBD-mi3); and N-glycan modified versions of the clade 1a and 1b RBDs (Wuhan-Hu-1 SARS-2, RaTG13, SHC014, pang17, and WIV1) together with unmodified Rs4081, RmYN02, and Rf1 RBDs (mosaic-8gm RBD-mi3). For mosaic-8 mi3 nanoparticles, equivalent conjugation of each of the eight SpyTagged RBDs was verified as described by SEC and SDS-PAGE analysis of conjugations to make homotypic nanoparticles (*34*).

Conjugated RBD-mi3 particles were separated from free RBDs by SEC on a Superose 6 10/300 column (GE Healthcare) equilibrated with PBS (20 mM sodium phosphate pH 7.5, 150 mM NaCl) and fractions corresponding to conjugated RBD-mi3 and free RBD were identified by SDS-PAGE. Concentrations of conjugated mi3 particles were determined using a Bio-Rad Protein Assay and are reported based on RBD content.

RBD-mi3 nanoparticles were evaluated for binding to a human ACE2-Fc construct (*32*) and to human monoclonal antibodies that recognize known RBD epitopes by ELISA. Duplicate samples of 20 µL of a 2.5 µg/mL solution of a purified RBD-mi3 nanoparticle in 0.1 M NaHCO_3_ pH 9.8 was coated onto Nunc® MaxiSorp™ 384-well plates (Sigma) and incubated overnight at 4°C. After blocking with 3% bovine serum albumin (BSA) in TBS containing 0.1% Tween 20 (TBS-T) for 1 hr at room temperature, plates were washed with TBS-T, and purified hACE2-Fc or human IgG (50 µg/mL with 8 4-fold serial dilutions in TBS-T/3% BSA) was added to plates for 3 hr at room temperature. Plates were then washed again for 1 hr at room temperature, and a 1:100,000 dilution of secondary HRP-conjugated goat anti-human IgG (Abcam) was added. SuperSignal™ ELISA Femto Maximum Sensitivity Substrate (ThermoFisher) was added to plates following manufacturer instructions, and plates were read at 425 nm. For the homotypic SARS-2 Beta ELISA shown in fig. S2F, 50 µL of a 2.5 µg/mL solution of a purified RBD-mi3 nanoparticle in 0.1 M NaHCO_3_ pH 9.8 was coated onto Corning® 96 well plates (Sigma) and incubated overnight at 4°C. After blocking with 3% bovine serum albumin (BSA) in TBS containing 0.1% Tween 20 (TBS-T) for 1 hr at room temperature, plates were washed with TBS-T, and purified hACE2-Fc or human IgG (50 µg/mL with 8 serial dilutions 4-fold in TBS-T/3% BSA) was added to plates for 3 hr at room temperature. Plates were then washed again for 1 hr at room temperature, and a 1:10,000 dilution of secondary HRP-conjugated goat anti-human IgG (Abcam) was added. 1-step Ultra TMB-ELISA (ThermoFisher) was added to plates following manufacturer instructions, and plates were read at 450 nm.

Prior to shipping for immunization and challenge studies, aliquots of conjugated RBD-mi3 nanoparticles were frozen in liquid nitrogen and then lyophilized (*35*) in PBS pH 7.4 using a Labconco CentriVap Benchtop Concentrator at -4°C. For immunizations, distilled water was added to rehydrate to a concentration of 1 mg/mL for a working stock, and the solution was gently pipetted and then spun at 20,000 x g for 10 minutes to remove aggregates.

### DLS and EM characterizations of RBD-mi3 nanoparticles

DLS was used to evaluate the hydrodynamic radii of conjugated nanoparticles. Lyophilized nanoparticles were rehydrated as described above. Sample sizes of 100 µL were loaded into a disposable cuvette, and DLS measurements were performed on a DynaPro® NanoStar^TM^ (Wyatt Technology) using settings suggested by the manufacturer. A fit of the second order autocorrelation function to a globular protein model was used to derive the hydrodynamic radius and plotted on Graphpad Prism 9.3.1.

Mosaic-8b RBD-mi3 and homotypic SARS-2 RBD-mi3 were compared by negative-stain EM. Ultrathin, holey carbon-coated, 400 mesh Cu grids (Ted Pella, Inc.) were glow discharged (60 s at 15 mA), and a 3 μL aliquot of SEC-purified RBD-mi3 nanoparticles were diluted to ∼40-100 ug/mL and applied to the grids for 60 s, Grids were then negatively stained with 2% (w/v) uranyl acetate for 30 s. Images were collected with a 120 keV FEI Tecnai T12 transmission electron microscope at 42,000x magnification.

### K18-hACE2 mice

The Institutional Animal Care and Use Committee at Rocky Mountain Laboratories provided animal study approvals, which were conducted in an Association for Assessment and Accreditation of Laboratory Animal Care-accredited facility, following the basic principles and guidelines in the *Guide for the Care and Use of Laboratory Animals* eighth edition, the Animal Welfare Act, U.S. Department of Agriculture, and the U.S. Public Health Service Policy on Humane Care and Use of Laboratory Animals.

Animals were kept in climate-controlled rooms with a fixed light/dark cycle (12 hours/12 hours). Mice were cohoused in rodent cages, fed a commercial rodent chow with *ad libitum* water, and monitored at least once daily. The Institutional Biosafety Committee (IBC)–approved work with infectious SARS-1 and SARS-2 viruses was conducted under biosafety level 3 (BSL3) conditions. All sample inactivation was performed according to IBC-approved standard operating procedures for removal of specimens from high containment.

### Cells and virus for K18-hACE2 mouse studies

Virus propagation was performed in VeroE6 cells in DMEM containing 2% FBS, 1 mM L-glutamine, penicillin (50 U/mL), and streptomycin (50 μg/mL (DMEM2). The consensus sequence of the virus stock (SARS-CoV-2 Beta, isolate hCoV-19/USA/MD-HP01542/2021) used for these experiments was identical to the initial sequence deposited on GISAID (EPI_ISL_890360), and no contaminants or additional mutations were detected. VeroE6 cells were maintained in DMEM supplemented with 10% fetal bovine serum, 1 mM L-glutamine, penicillin (50 U/mL), and streptomycin (50 μg/mL). VeroE6 cells were provided by R. Baric (University of North Carolina at Chapel Hill). Mycoplasma testing is performed at monthly intervals, and no mycoplasma was detected.

### Vaccination and infection of K18-hACE2 mice

K18-hACE2 mice (4 to 6 weeks old) were vaccinated with 2 x 50 μL of 5 µg (RBD equivalents)/(11.4 µg of total RBD-mi3) of RBD-mi3 or 5 µg unconjugated mi3 adjuvanted with Addavax 1:1 (1:1 v/v) intramuscularly at day 0 and day 28 (and challenged 28 days post the second immunization). Fourteen days before virus challenge, animals were bled via the submandibular vein. 10 animals per group were challenged with 30 μL of 10^5^ TCID_50_ SARS-2/human/USA/MD-HP01542/2021) or SARS-1 (Tor2) diluted in sterile Dulbecco’s modified Eagle’s medium (DMEM). Weight was recorded daily. Six mice per group were observed for survival up to 28 days post challenge or until they reached end-point criteria. End-point criteria were as follows: labored breathing or ambulatory difficulties or weight loss exceeding 20%. Four animals per group were euthanized on day 4 post challenge to collect oropharyngeal swabs and lung tissue for virology and histology analysis.

### Virus titration after K18-hACE2 mouse challenge

Lung tissue sections were weighed and homogenized in 750 μL of DMEM. Virus titrations were performed by end point titration in VeroE6 cells expressing transmembrane protease serine 2 (TMPRSS-2) and human ACE2 (BEI resources, NR-54970), which were inoculated with 10-fold serial dilutions of virus swab medium or tissue homogenates in 96-well plates. When titrating tissue homogenate, cells were washed with PBS and 100 μL of DMEM2. Cells were incubated at 37°C and 5% CO_2_, and cytopathic effect (CPE) was assessed 6 days later.

### RNA extraction and quantitative RT-PCR

RNA was extracted from oropharyngeal swabs swabs using a QIAamp Viral RNA kit (Qiagen) according to the manufacturer’s instructions. Tissue was homogenized and extracted using the RNeasy kit (Qiagen) according to the manufacturer’s instructions. Viral gRNA- and sgRNA-specific assays (*47*) were used for the detection of viral RNA. The RT-PCR reaction (with 5 μL template viral RNA) was performed using the QuantStudio (Thermo Fisher Scientific) according to instructions of the manufacturer. Dilutions of SARS-2 with known genome copies were run in parallel to be used to generate the standard curves.

### Histopathology of K18-hACE2 mouse samples

Lungs were collected upon necropsy on day 4 post challenge and perfused with 10% neutral-buffered formalin. Fixation was done for at least 7 days. Tissues were placed in cassettes and processed with a Sakura VIP-6 Tissue Tek on a 12-hour automated schedule using a graded series of ethanol, xylene, and PureAffin. Embedded tissues were sectioned at 5µm and dried overnight at 42 degrees C prior to staining. Sections were stained with Harris hematoxylin (Cancer Diagnostics, no. SH3777), decolorized with 0.125% HCl/70% ethanol, blued in Pureview PH Blue (Cancer Diagnostics, no. 167020), counterstained with eosin 615 (Cancer Diagnostics, no. 16601), dehydrated, and mounted in Micromount (Leica, no. 3801731). An anti-SARS-2 nucleocapsid protein rabbit antiserum (generated by GenScript) was used at a 1:1000 dilution to detect specific anti–SARS-2 immunoreactivity using the Discovery ULTRA automated staining instrument (Roche Tissue Diagnostics) with a Discovery ChromoMap DAB (Ventana Medical Systems) kit. All slides were examined by a board-certified veterinary anatomic pathologist who was blinded to study group allocations. Scoring was done as follows. H & E; no lesions = 0; less than 1% = 0.5; minimal (1-10%) = 1; mild (11-25%) = 2; moderate (26-50%) = 3; marked (51-75%) = 4; severe (76-100%) = 5. IHC attachment; none = 0; less than 1% = 0.5; rare/few (1-10%) = 1; scattered (11-25%) = 2; moderate (26-50%) = 3; numerous (51-75%) = 4; diffuse (76-100%) = 5. Histopathology report is summarized in Data S1.

### BIOQUAL Ethics Statement and Animal Exposure

Rhesus macaques were housed and cared for at BIOQUAL, Inc., Rockville, MD. The study was performed under a BIOQUAL-approved IACUC protocol (no. 21-092P), in strict accordance with the recommendations in the Guide for the Care and Use of Laboratory Animals of the NIH, and in accordance with BIOQUAL standard operating procedures. BIOQUAL is fully accredited by the Association for Assessment and Accreditation of Laboratory Animal Care (AAALAC) and through OLAW, assurance number A-3086. All animal procedures were done under anesthesia to minimize pain and distress, in accordance with the recommendations of the Weatherall report ‘The use of non-human primates in research.’ Teklad 5038 primate diet was provided once daily according to macaque size and weight. The diet was supplemented daily with fresh fruit and vegetables. Fresh water was given *ad libitum*.

### Vaccination of NHPs

The study included 16 rhesus macaques (Macaca mulatta), 8 of which were immunized with mosaic-8b RBD-mi3 (n = 8), and 8 of which served as unimmunized controls for SARS-2 and SARS-1 challenges. Four immunized and four unimmunized control NHPs were challenged with SARS-2, and four immunized and four unimmunized control NHPs were challenged with SARS-1. Due to a shortage of available NHPs, we could not compare mosaic-8b RBD-mi3 and homotypic SARS-2 Beta RBD-mi3 immunizations in this study. Macaques were 3-5 years old and ranged from 3.2 to 5.1 kg in body weight. Male and female macaques per group were balanced. Studies were performed unblinded. Macaques were evaluated by BIOQUAL veterinary staff before, during, and after immunizations.

NHPs were immunized intramuscularly with 25 μg (calculated based on RBDs; 56.8 µg of total RBD-mi3) of mosaic-8b RBD-mi3 adjuvanted with VAC20 (2% aluminum hydroxide wet gel, Al_2_O_3_) (alum) (Prime and Boost 1) (kind gift of Francis Laurent and Ruben Caputo, SPI Pharma) and subsequently with MF59 adjuvant (EmulsiPan, a squalene-based oil in water emulsion adjuvant (*50*) (kind gift of Harshet Jain, Panacea Biotec) for Boost 2. Each macaque received 0.5 mL into the right forelimb.

### SARS-2 and SARS-1 intranasal and intratracheal NHP challenges

All macaques were challenged at week 11 (3 weeks after last vaccination) through combined intratracheal (1.0 mL) and intranasal (0.5 mL per nostril) inoculation with an infectious dose of 10^5 TCID_50_ of SARS-2 B.1.617.2 (Delta, BEI NR-55612) or SARS-1 (Urbani). Virus was stored at −80 °C before use, thawed by hand and placed immediately on wet ice. Stock was diluted to 5 × 10^4 TCID_50_ mL^−1^ in PBS and vortexed gently for 5 s before inoculation. Nasal swabs, BAL, plasma, and serum samples were collected seven days before and two and four days after challenge. Protection from SARS-2 and SARS-1 infection was determined by quantitative infectious viral load assay (TCID_50_), and for SARS-2, also by RT-PCR of subgenomic N RNA (N sgRNA) as described above except that amplification was done use the Applied Biosystems 7500 Sequence detector.

### TCID_50_ and SARS-2 and SARS-1 virus PRNT_50_ assays in NHP samples

PRNT_50_ (50% plaque reduction neutralization test) assays for NHP samples were performed in a biosafety level 3 facility at BIOQUAL, Inc. (Rockville, MD). The TCID_50_ assay was conducted by addition of 10-fold graded dilutions of samples to Vero/TMPRSS2 cell monolayers. Serial dilutions were performed in cell culture wells in quadruplicates. Positive (virus stock of known infectious titer in the assay) and negative (medium only) control wells were included in each assay set-up. The plates were incubated at 37°C, 5.0% CO_2_ for 4 days. The cell monolayers were visually inspected for CPE, i.e., complete destruction of the monolayer. TCID_50_ values was calculated using the Reed-Muench formula (*85*). For samples that had less than 3 CPE positive wells, the TCID_50_ could not be calculated using the Reed-Muench formula, and these samples were assigned a titer of below the limit of detection (i.e., <2.7 log10 TCID_50_/mL). For acceptable assay performance, the TCID_50_ value of the positive control tested within 2-fold of the expected value.

To measure neutralization activity, sera from each NHP were diluted to 1:10 followed by a 3-fold serial dilution. Diluted samples were then incubated with ∼30 plaque-forming units of wild-type SARS-2 USA-WA1/2020 (BEI NR-52281), B.1.351 (Beta, 501Y.V2.HV, NR-54974), or B.1.617.2 (Delta, BEI NR-55612) variants, in an equal volume of culture medium for 1 hour at 37°C. The serum-virus mixtures were added to a monolayer of confluent Vero E6 cells and incubated for one hour at 37°C in 5% CO_2_. Each well was then overlaid with culture medium containing 0.5% methylcellulose and incubated for 3 days at 37°C in 5% CO_2_. The plates were then fixed with methanol at -20°C for 30 minutes and stained with 0.2% crystal violet for 30 min at room temperature. PRNT_50_ were estimated by determining the dilution at which plaques were reduced by 50% with respect to viral control.

### Mouse and NHP serum ELISAs

20 µL of a 2.5 µg/mL solution of an affinity purified His-tagged RBD in 0.1 M NaHCO_3_ pH 9.8 was coated onto Nunc® MaxiSorp™ 384-well plates (Sigma) and incubated overnight at 4°C. After blocking with 3% bovine serum albumin (BSA) in TBS containing 0.1% Tween 20 (TBS-T) for 1 hr at room temperature, plates were washed with TBS-T, and mouse or NHP serum diluted 1:100 and then serially diluted by 4-fold with TBS-T/3% BSA was added to the plates for 3 hr at room temperature. Plates were then washed again for 1 hr at room temperature, and a 1:50,000 dilution of secondary HRP-conjugated goat anti-mouse IgG (Abcam) was added. SuperSignal™ ELISA Femto Maximum Sensitivity Substrate (ThermoFisher) was added to plates following manufacturer’s instructions, and plates were read at 425 nm. Curves were plotted and analyzed to obtain midpoint titers (EC_50_ values) using Graphpad Prism 9.3.1 (Graphpad Softwatre, San Diego, CA) assuming a one-site binding model with a Hill coefficient. Titer differences were evaluated for statistical significance between groups using ANOVA test followed by Tukey’s multiple comparison post hoc test calculated using Graphpad Prism 9.3.1.

### Mouse and NHP serum pseudovirus neutralization assays

Lentiviral-based SARS-2 variants (Wuhan-Hu-1, Beta, Delta, Omicron BA.1), SARS-1, WIV1, SHC014, and BtKY72 K493Y/T498W (*13*) (kind gift of Alexandra Walls and David Veesler, University of Washington) pseudoviruses were prepared as described (*20, 86*) using genes encoding S protein sequences lacking C-terminal residues in the cytoplasmic tail: 21 amino acid deletions for SARS-2 variants, WIV1, SHC014, and BtKY72 and a 19 amino acid deletion for SARS-CoV. For neutralization assays, three-fold serially diluted sera from immunized mice or NHPs were incubated with a pseudovirus for 1 hour at 37°C, then the serum/virus mixture was added to 293T_ACE2_ target cells and incubated for 48 hours at 37°C. Media was removed, cells were lysed with Britelite Plus reagent (Perkin Elmer), and luciferase activity was measured as relative luminesce units (RLUs). Relative RLUs were normalized to RLUs from cells infected with pseudotyped virus in the absence of antiserum. Half-maximal inhibitory dilutions (ID_50_ values) were derived using 4-parameter nonlinear regression in AntibodyDatabase (*87*). Statistical significance of titer differences between groups were evaluated using ANOVA test followed by Tukey’s multiple comparison post hoc test of ID_50_s converted to log10 scale using Graphpad Prism 9.3.1.

### Mouse serum samples for RBD epitope mapping

Animal procedures and experiments were performed at Labcorp Drug Development (formerly Covance, Inc.) according to protocols approved by the IACUC to obtain serum samples for RBD epitope mapping experiments. Immunizations of mosaic-8b or homotypic SARS-2 Beta (5 µg each based on RBD content, 11.4 µg of total RBD-mi3) in 100 µL of 50% v/v AddaVax^TM^ adjuvant (InvivoGen) were done using intramuscular (IM) injections of 7-8-week-old female BALB/c mice (Envigo) (8 animals per cohort). Animals were boosted 3 weeks after the prime with the same quantity of antigen in adjuvant. Animals were bled under anesthesia approximately every 2 weeks via orbital sinus and then euthanized 7 weeks after the prime (Day 49) after blood collection from the jugular vein. Blood samples were stored at room temperature in serum separator tubes (BD Microtainer) to allow clotting. Serum was then harvested into microtubes (Mikro-Schraubrohre) and stored at -80°C until use.

### RBD sequencing library construction and SARS-2 enrichment

To construct sequencing libraries for RBD epitope mapping of mouse sera, 25 μl of ds-cDNA was brought to a final volume of 53 μL in elution buffer (Agilent Technologies) and sheared on a Covaris LE220 (Covaris) to generate an average size of 180 to 220 base pairs (bp). The following settings were used: peak incident power, 450 W; duty factor, 15%; cycles per burst, 1000; and time, 300 s. The Kapa HyperPrep kit was used to prepare libraries from 50 μL of each sheared cDNA sample following modifications of the Kapa HyperPrep kit (version 8.20) and SeqCap EZ HyperCap Workflow (version 2.3) user guides (Roche Sequencing Solutions Inc.). Adapter ligation was performed for 1 hour at 20°C using the Kapa Unique-Dual Indexed Adapters diluted to 1.5 μM concentration (Roche Sequencing Solutions Inc.). After ligation, samples were purified with AmPure XP beads (Beckman Coulter) and subjected to double-sided size selection as specified in the SeqCap EZ HyperCap Workflow User’s Guide. Precapture polymerase chain reaction (PCR) amplification was performed using 12 cycles, followed by purification using AmPure XP beads. Purified libraries were assessed for quality on the Bioanalyzer 2100 using the High-Sensitivity DNA chip assay (Agilent Technologies). Quantification of pre-capture libraries was performed using the Qubit dsDNA HS Assay kit and the Qubit 3.0 fluorometer following the manufacturer’s instructions (Thermo Fisher Scientific).

The myBaits Expert Virus bait library was used to enrich samples for SARS-2 according to the myBaits Hybridization Capture for Targeted NGS (version 4.01) protocol. Briefly, libraries were sorted according to estimated genome copies and pooled to create a combined mass of 2 μg for each capture reaction. Depending on estimated genome copies, two to six libraries were pooled for each capture reaction. Capture hybridizations were performed for 16 to 19 hours at 65°C and subjected to 8 to 14 PCR cycles after enrichment. SARS-2–enriched libraries were purified and quantified using the Kapa Library Quant Universal quantitative PCR mix in accordance with the manufacturer’s instructions. Libraries were diluted to a final working concentration of 1 to 2 nM, titrated to 20 pM, and sequenced as 2 × 150 bp reads on the MiSeq sequencing instrument using the MiSeq Micro kit version 2 (Illumina).

### Sorting of yeast libraries to identify mutations that reduced binding by polyclonal antisera

Plasma mapping experiments were performed in biological duplicate using the independent mutant RBD libraries as previously described (*44*). Prior to the yeast-display deep mutational scanning experiments, 100 µL of each serum sample was heat-inactivated at 56°C for 30 min and twice-depleted of nonspecific yeast-binding antibodies by incubating with 50 OD units of AWY101 yeast containing an empty vector (*54*). Mutant yeast libraries that were pre-sorted for RBD expression and ACE2 binding (*54*) were induced to express RBD in galactose-containing synthetic defined medium with casamino acids (6.7g/L Yeast Nitrogen Base, 5.0g/L Casamino acids, 1.065 g/L MES acid, and 2% w/v galactose + 0.1% w/v dextrose). 16–18 hours post-induction, cells were washed and incubated with plasma at a range of dilutions for 1 hour at room temperature with gentle agitation. For each plasma, we chose a sub-saturating dilution such that the amount of fluorescent signal due to plasma antibody binding to RBD was approximately equal across samples. The exact dilution used for each plasma is given in fig. S5B. The libraries were washed and secondarily labeled for 1 hour with 1:100 fluorescein isothiocyanate-conjugated anti-MYC antibody (Immunology Consultants Lab, CYMC-45F) to label for RBD expression and 1:200 Alexa Fluor-647-conjugated goat anti-human-IgG Fc-gamma (Jackson ImmunoResearch 109-135-098) to label for bound NHP antibodies or Alexa Fluor-647-conjugated goat anti-mouse-IgG Fc-gamma (Jackson ImmunoResearch 115-605-008) to label for bound mouse antibodies. A flow cytometric selection gate was drawn to capture RBD mutants with reduced antibody binding for their degree of RBD expression (fig. S5C). For each sample, 7.5 x 10^6^ to 1.1 x 10^7^ cells were processed on the BD FACSAria II cell sorter (fig. S5C). Antibody-escaped cells were grown overnight in synthetic defined medium with casamino acids (6.7g/L Yeast Nitrogen Base, 5.0g/L Casamino acids, 1.065 g/L MES acid, and 2% w/v dextrose + 100 U/mL penicillin + 100 µg/mL streptomycin) to expand cells prior to plasmid extraction.

### DNA extraction and Illumina sequencing

Plasmid samples were prepared from 30 optical density (OD) units (1.6e8 colony forming units (cfus)) of pre-selection yeast populations and approximately 5 OD units (∼3.2e7 cfus) of overnight cultures of plasma-escaped cells (Zymoprep Yeast Plasmid Miniprep II) as previously described (*53, 88*). The 16-nucleotide barcode sequences identifying each RBD variant were amplified by polymerase chain reaction (PCR) and prepared for Illumina sequencing as described (*53, 88*). Barcodes were sequenced on an Illumina HiSeq 2500 with 50 bp single-end reads. Raw sequencing data are available on the NCBI SRA under BioProject PRJNA770094, BioSample SAMN26315988.

### Analysis of deep sequencing data to compute each mutation’s escape fraction

Escape fractions were computed essentially as described in (*53*) and exactly as described in (*54*). We used the dms_variants package (https://jbloomlab.github.io/dms_variants/, version 0.8.10) to process Illumina sequences into counts of each barcoded RBD variant in each pre-selection and antibody-escape population. We computed the escape fraction for each barcoded variant using the deep sequencing counts for each variant in the original and plasma-escape populations and the total fraction of the library that escaped antibody binding via the formula in (*54*). These escape fractions represent the estimated fraction of cells expressing that specific variant that falls in the escape bin, such that a value of 0 means the variant is always bound by plasma and a value of 1 means that it always escapes plasma binding.

We then applied a computational filter to remove variants with >1 amino-acid mutation, low sequencing counts, or highly deleterious mutations that might escape antibody binding due to poor RBD expression or folding as described (*54*). The reported antibody-escape scores are the average across duplicate libraries; these scores are also in Data S2. Correlations in final single-mutant escape scores are shown in fig. S5D. Full documentation of the computational analysis is at https://github.com/jbloomlab/SARS-CoV-2-RBD_Beta_mosaic_np_vaccine.

### Data visualization

The static logo plot visualizations of the escape maps in the paper figures were created using the dmslogo package (https://jbloomlab.github.io/dmslogo, version 0.6.2) and in all cases the height of each letter indicates the escape fraction for that amino-acid mutation calculated as described above. For the mouse sera, the static logo plots feature any site where for >=1 serum, the site-total antibody escape was >10x the median across all sites and at least 10% the maximum of any site. Due to the relative breadth of the NHP sera, a more sensitive threshold for displaying sites on logo plots was used: we include any site where the site-total antibody escape is >5x the median across all sites and at least 5% maximum of any sites. This resulted in sites 383, 386, and 500. Thus, sites 346, 352, 357, 369, 378, 384, 385, 390, 396, 408, 462, 468, 477, 478, 484, 485, 486, and 501 were also added to the logo plots to facilitate comparison to the mouse sera. For each sample, the y-axis is scaled to be the greatest of (a) the maximum site-wise escape metric observed for that sample, or (b) 20x the median site-wise escape fraction observed across all sites for that plasma. The code that generates these logo plot visualizations is available at https://github.com/jbloomlab/SARS-CoV-2-RBD_Beta_mosaic_np_vaccine/blob/main/results/summary/escape_profiles.md. In many of the visualizations, the RBD sites are categorized by epitope region (23) and colored accordingly. Specifically, we define the class 1 epitope as residues 403+405+406+417+420+421+453+455– 460+473–478+486+487+489+503+504, the class 2 epitope as residues 472+479+483–485+490–495, the class 3 epitope to be residues 341+345+346+354–357+396+437-452466–468+496+498–501, and the class 4 epitope as residues 365–390+408+462.

For the static structural visualizations in figures, the Beta RBD surface (PDB 7LYQ) was colored by the site-wise escape metric at each site, with white indicating no escape and red scaled to be the same maximum used to scale the y-axis in the logo plot escape maps, determined as described above. We created interactive structure-based visualizations of the escape maps using dms-view (*89*) that are available at https://jbloomlab.github.io/SARS-CoV-2-RBD_Beta_mosaic_np_vaccine/.

---

**Figure S1.**
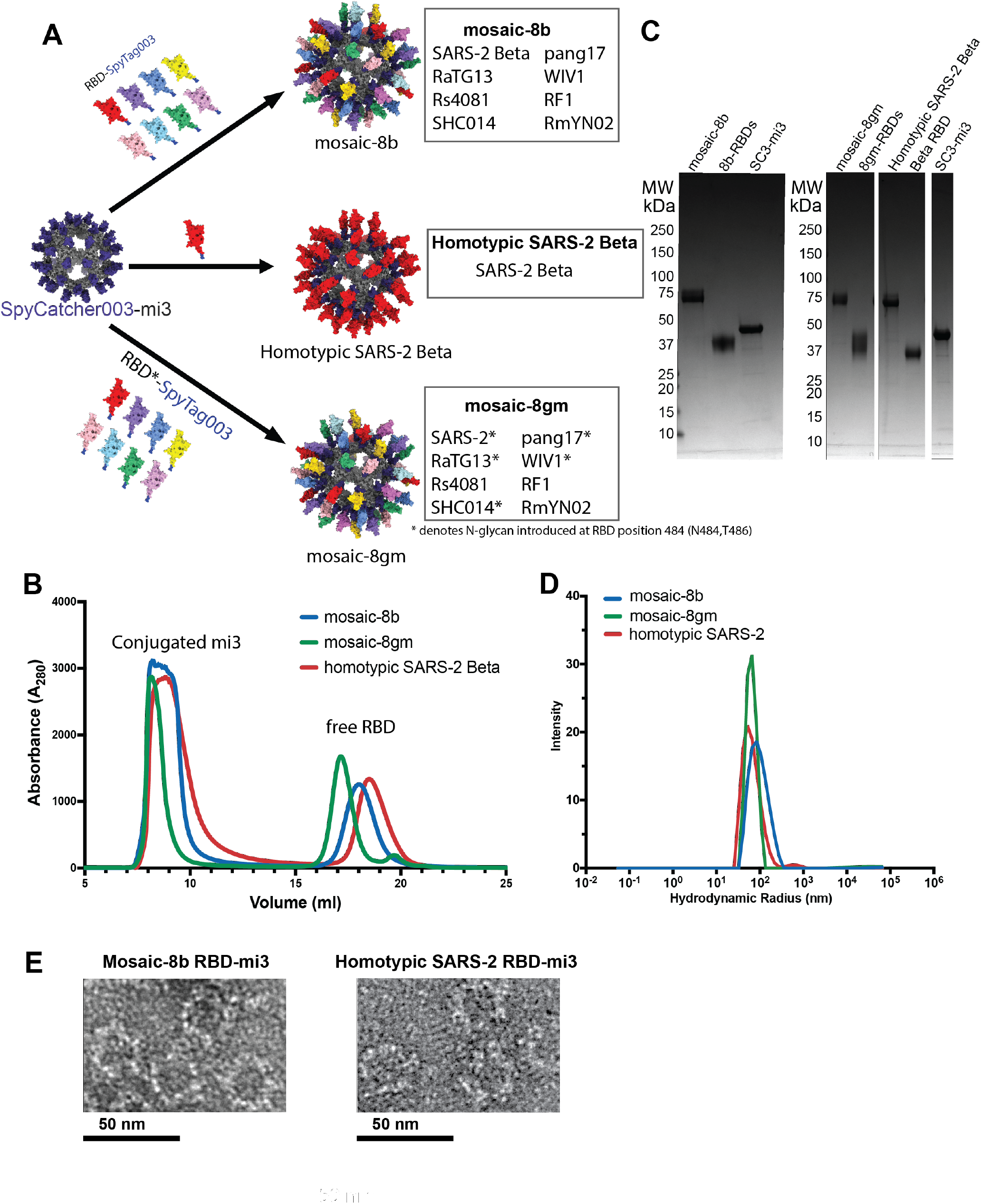
Preparation of RBD-mi3 nanoparticles. (**A**) Schematic for construction of mosaic-8b, mosaic-8gm, and homotypic SARS-2 Beta RBD-mi3 nanoparticles. (**B**) Superose 6 10/300 SEC profile after RBD conjugations to mi3 (2-fold molar excess of RBD to mi3 subunit) showing peaks for RBD-mi3 nanoparticles and free RBD(s). (**C**) SDS-PAGE (Coomassie staining) of RBD-coupled nanoparticles, free RBDs, and free SpyCatcher003-mi3 particles (SC3-mi3). (**D**) Dynamic light scattering (DLS) measurements for RBD-coupled nanoparticles. (**E**) Negative-stain EM images of RBD-coupled mi3 nanoparticles.

**Figure S2.**
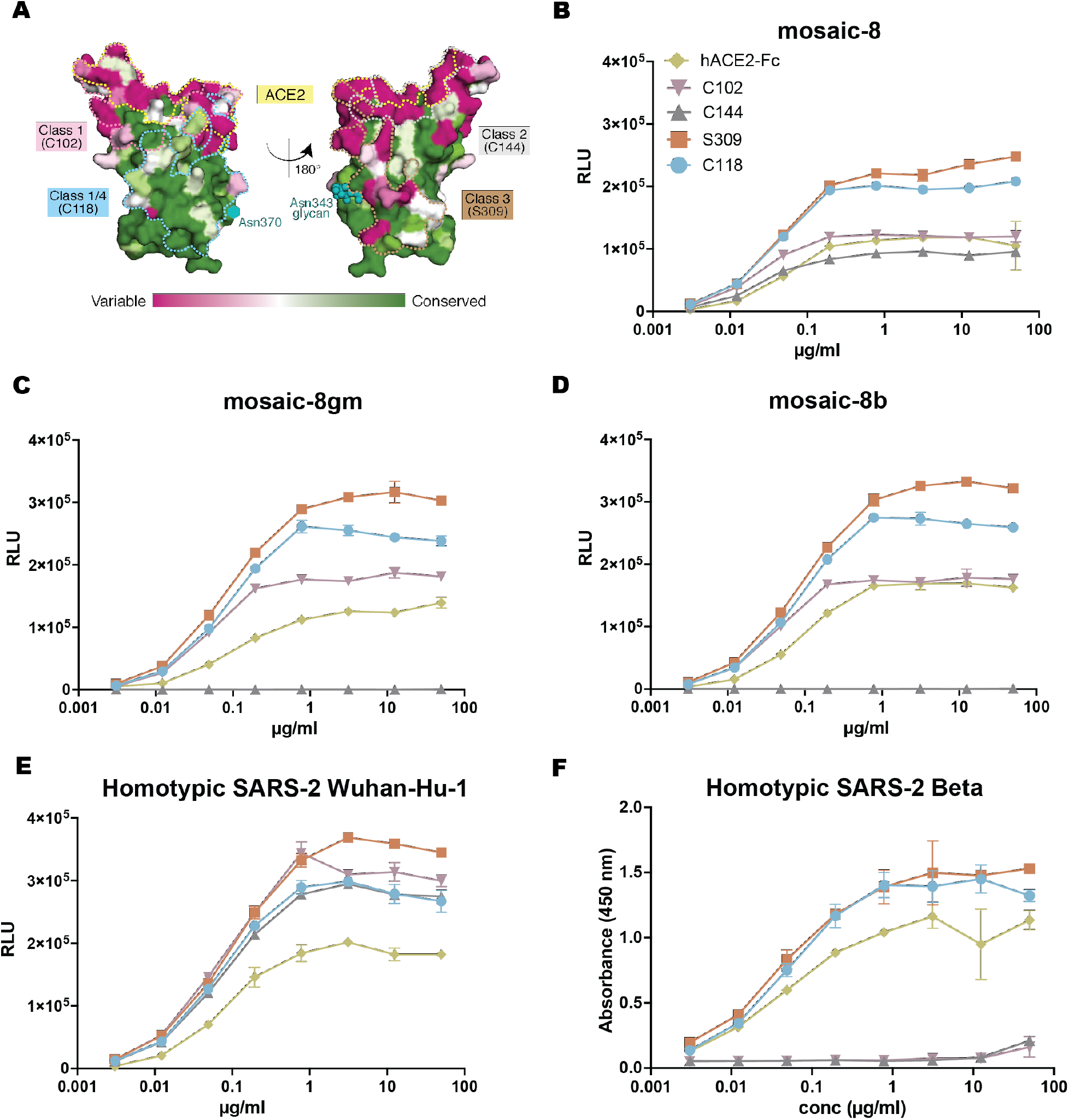
Binding characteristics of mosaic and homotypic RBD-mi3 nanoparticles. (**A**) Sequence conservation of the 16 sarbecovirus RBDs in Fig. 1D calculated by the ConSurf Database (*79*) shown on two views of an RBD surface (PDB 7BZ5). The ACE2 binding footprint (PDB 6M0J) is outlined by yellow dots. Epitopes of representative monoclonal antibodies used in binding experiments are outlined in dots of the indicated colors using information from structures of Fabs bound to RBD or S trimer (C118: PDB 7RKS, S309: PDB 7JX3; C144: PDB 7K90, C102: PDB 7K8M). The N-linked glycan attached to RBD residue 343 is indicated by teal spheres, and the potential N-linked glycosylation site at position 370 in RBDs derived from sarbecoviruses other than SARS-2 is indicated by a teal circle. (**B-F**) ELISAs to assess binding of the hACE2-Fc and the indicated monoclonal antibodies to RBD-mi3 nanoparticles. Nanoparticles were immobilized on an ELISA plate, incubated with the indicated monoclonal antibody or hACE2-Fc, and binding was detected using a labeled anti-human IgG secondary antibody. Data points are presented as the mean and standard deviation of duplicate measurements. Some error bars are too small to be distinguished from data points. RLU = relative luminescence units. (**B**) Binding to mosaic-8 RBD-mi3 (Wuhan-Hu-1 SARS-2 RBD plus seven animal sarbecovirus RBDs as previously described (*34*) and in fig. S1A). (**C**) Binding to mosaic-8gm RBD-mi3 (mosaic-8 with a Wuhan-Hu-1 SARS-2 RBD plus the seven animal sarbecovirus RBDs in which N-linked glycosylation site sequons at RBD position 484 were introduced in the clade 1a and 1b RBDs to occlude class 1 and 2 RBD epitopes). (**D**) Binding to mosaic-8b RBD-mi3 (SARS-2 Beta RBD plus the seven animal sarbecovirus RBDs in fig. S1A). (**E**) Binding to homotypic SARS-2 Wuhan Hu-1 RBD-mi3 (as previously described (*34*)). (**F**) Binding to homotypic SARS-2 Beta RBD-mi3.

**Figure S3.**
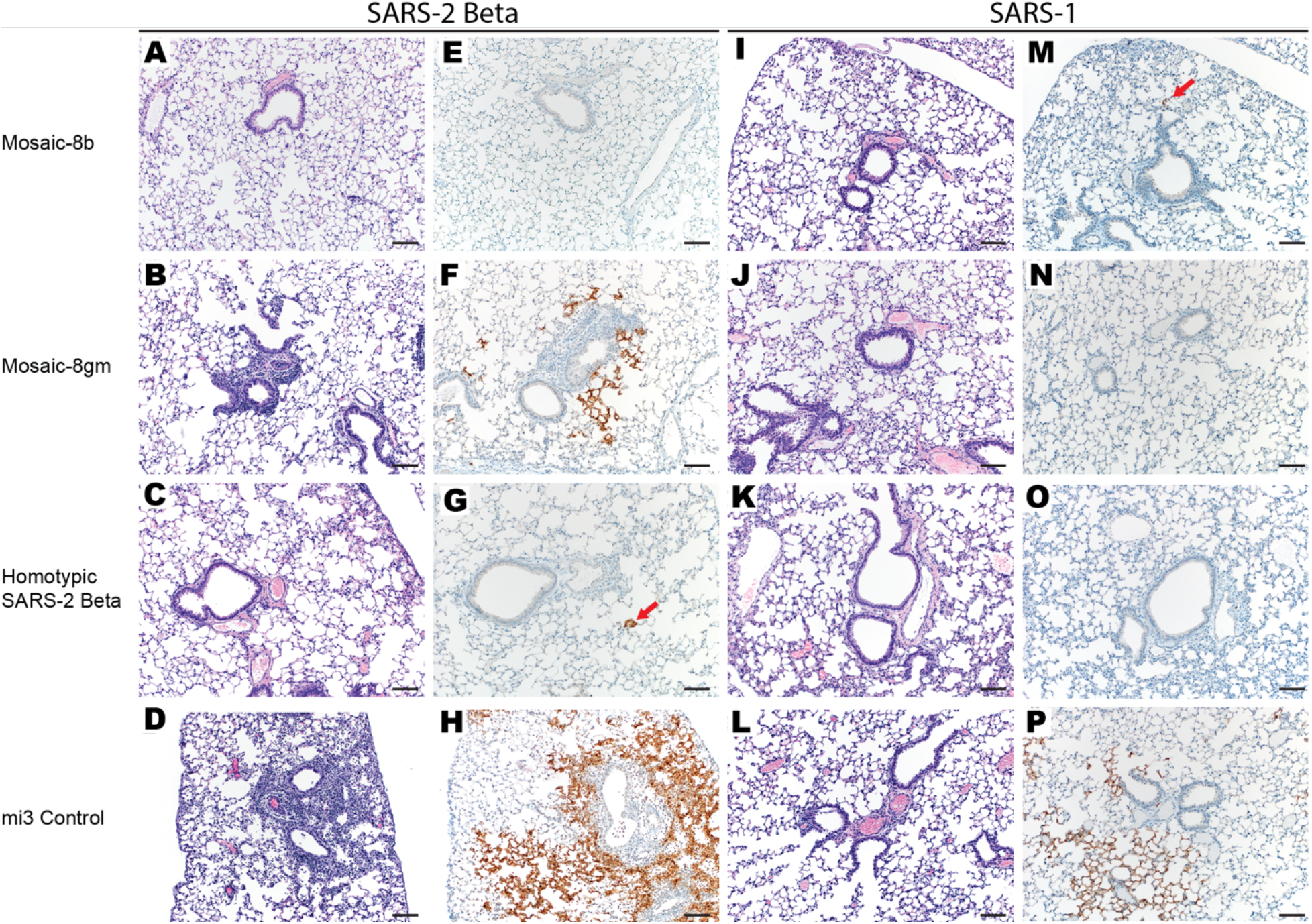
Lung pathology is reduced in mosaic-8b immunized mice challenged with either SARS-2 or SARS-1. Images taken at 100x magnification. Scale bar = 100µm. Red arrows = immunoreactivity in panels G and M. (**A-D)** Hematoxylin and eosin (H&E) stained lung tissue sections from animals vaccinated with either mosaic-8b, mosaic-8gm, homotypic SARS-2 Beta, or unconjugated mi3 and challenged with SARS-2 Beta (minimal-mild peribronchial inflammation in panels B and D). **(E-H)** Immunohistochemistry (IHC) staining for SARS-CoV-2 N protein antigen from animals vaccinated with either mosaic-8b, mosaic-8gm, homotypic SARS-2 Beta, or mi3 and challenged with SARS-2 Beta. **(I-L)** H&E stained lung tissue sections from animals vaccinated with either mosaic-8b, mosaic-8gm, homotypic SARS-2 Beta, or mi3 and challenged with SARS-1. **(M-P)** Immunohistochemistry staining for SARS-CoV-2 N protein antigen from animals vaccinated with either mosaic-8b, mosaic-8gm, homotypic SARS-2 Beta, or mi3 and challenged with SARS-1.

**Figure S4.**
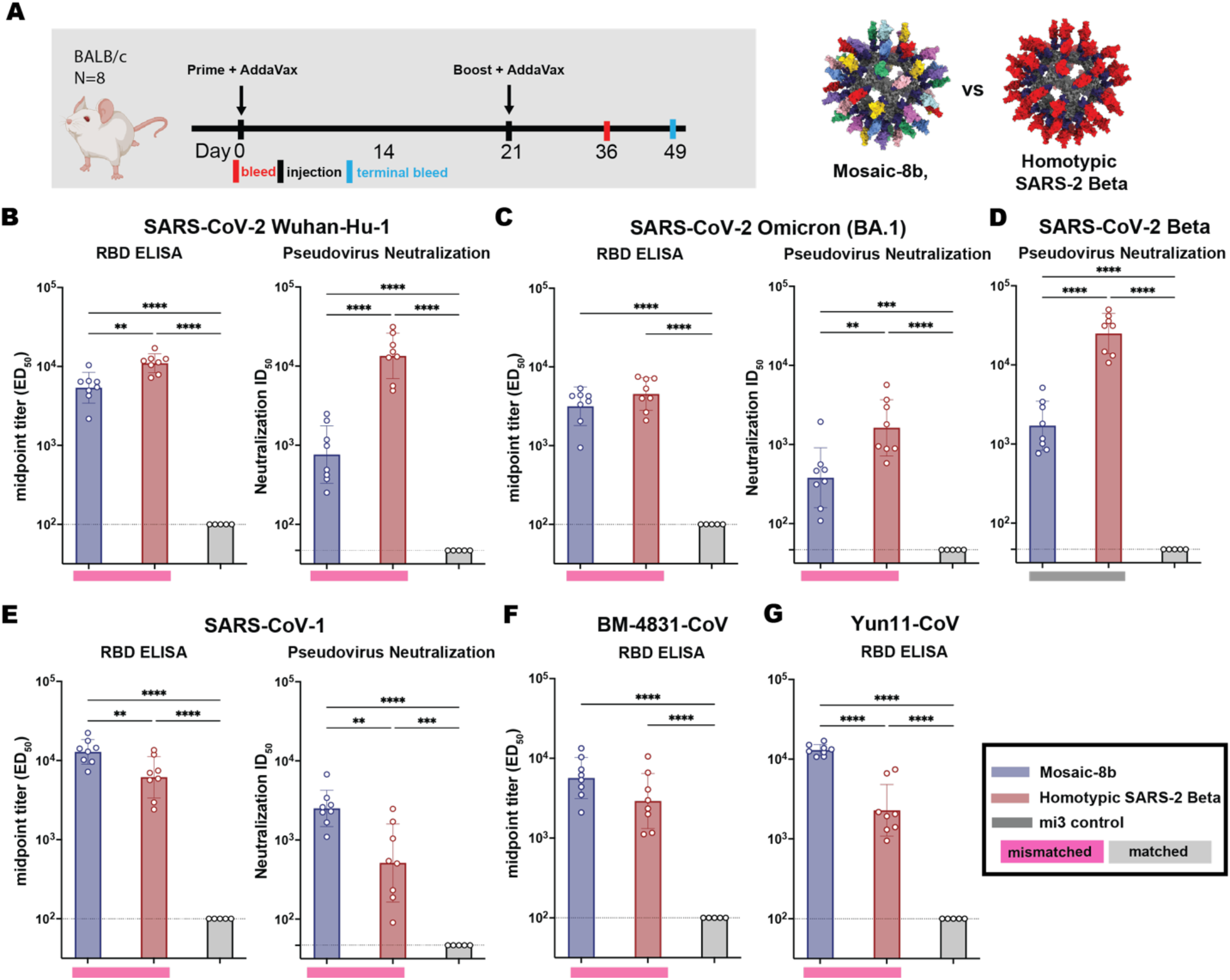
Mosaic-8b and homotypic SARS-2 Beta RBD-mi3 immunizations elicit binding and neutralizing antibodies in BALB/c mice. (**A**) Left: Immunization schedule. BALB/c mice were immunized with either mosaic-8 or homotypic SARS-2 Beta RBD-mi3. Right: Structural models of mosaic-8 and homotypic RBD-mi3 nanoparticles constructed using PDB 7SC1 (RBD), PDB 4MLI (SpyCatcher), and PDB 7B3Y (mi3). (**B-G**) ELISA and neutralization data for antisera (taken 4 weeks post boost) from individual mice (open circles) presented as the mean (bars) and standard deviation (horizontal lines). ELISA results are shown as midpoint titers (EC_50_ values); neutralization results are shown as half-maximal inhibitory dilutions (ID_50_ values). Dashed horizontal lines correspond to the background values representing the limit of detection. Significant differences between cohorts linked by horizontal lines are indicated by asterisks: p<0.05 = *, p<0.01 = **, p<0.001 = ***, p<0.0001 = ****. Rectangles below ELISA and neutralization data indicate mismatched strains (pink; the RBD from that strain was not present on the nanoparticle) or matched strains (gray; the RBD was present on the nanoparticle).

**Figure S5.**
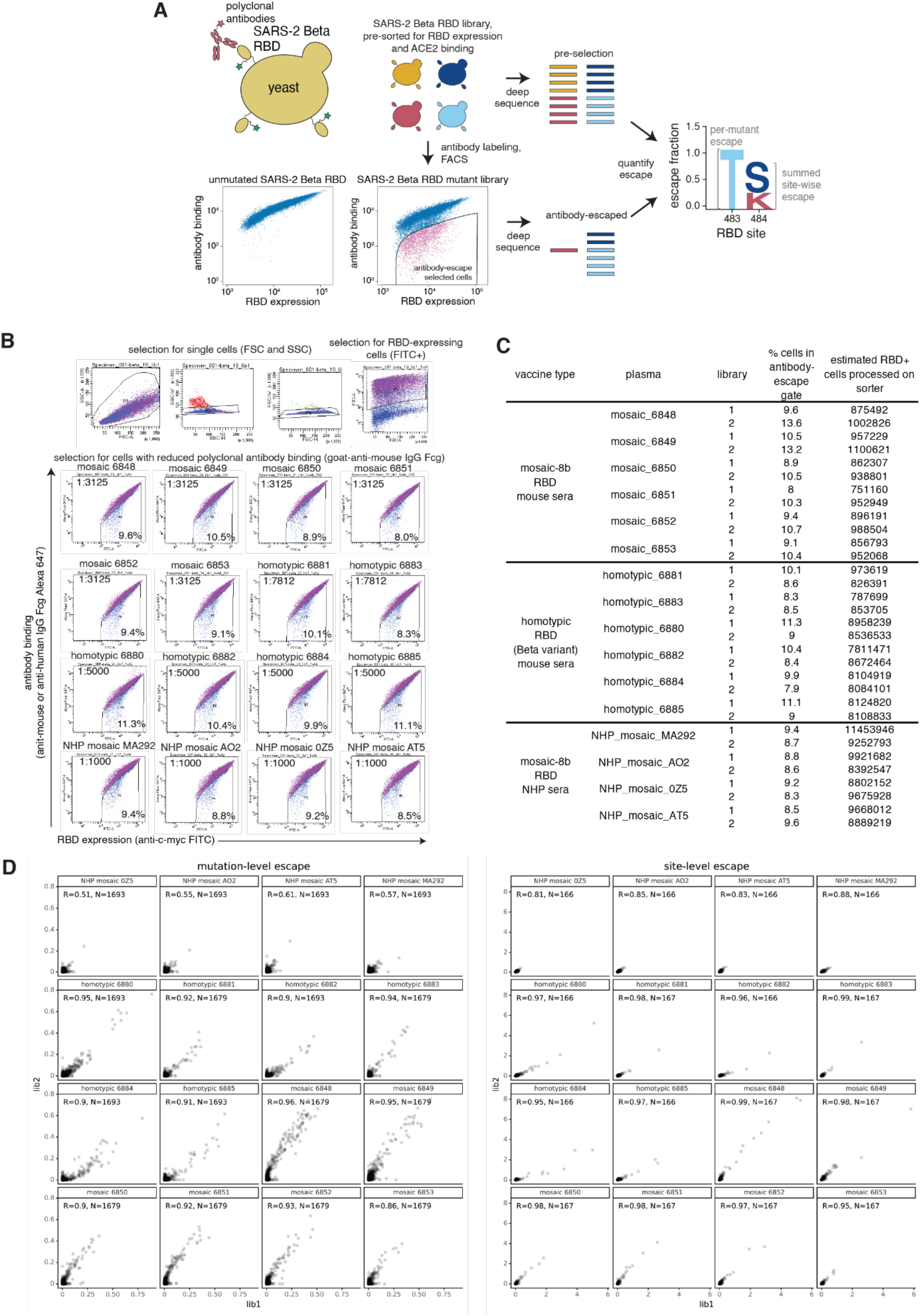
Comprehensive mapping of mutations that reduce binding of sera from immunized mice or NHPs to the SARS-2 Beta RBD. (**A**) Deep mutational scanning (*54*) was used to map mutations that reduced binding of polyclonal antibodies from immunized animals to the SARS-2 Beta RBD. A library of yeast containing nearly all possible mutations in the SARS-2 Beta RBD was incubated with sera from immunized mice or NHPs, and fluorescence-activated cell sorting (FACS) was used to enrich for cells expressing RBD (detected with a C-terminal Myc tag, green star) with reduced antibody binding, detected using an anti-mouse (for mouse sera) or anti-human (for NHP sera) IgG Fc-gamma secondary antibody. Deep sequencing was used to quantify the frequency of each mutation in the pre-selection and antibody-escape cell populations. We calculated each mutation’s “escape fraction,” the fraction of cells expressing RBD with that mutation that fell in the antibody-escape FACS bin (ranging from 0 to 1). The site-level escape metric is the sum of the escape fractions of all mutations at a site. (**B**) Top: Representative plots of nested FACS gating strategy used for all experiments to select for RBD+ single cells. Samples were gated by SSC-A versus FSC-A, SSC-W versus SSC-H, and FSC-W versus FSC-H) that also express RBD (FITC-A vs. FSC-A). Bottom: FACS gating strategy for one of two independent libraries to select cells expressing RBD mutants with reduced binding by polyclonal sera (cells in blue). Gates were set manually during sorting. Selection gates were set to capture cells that have a reduced amount of antibody binding for their degree of RBD expression. FACS scatter plots were qualitatively similar between the two libraries. SSC-A, side scatter-area; FSC-A, forward scatter-area; SSC-W, side scatter-width; SSC-H, side scatter-height; FSC-W, forward scatter-width; FSC-H, forward scatter height; FITC-A, fluorescein isothiocyanate-area. (**C**) The percent and number of RBD+ cells sorted into the antibody-escape gate for each library selected against each serum. (**D**) Mutation (top)- and site (bottom)-level correlations of escape scores between two independent biological replicate libraries. The complete antibody-escape scores are available in Data S2 and at https://github.com/jbloomlab/SARS-CoV-2-RBD_Beta_mosaic_np_vaccine/blob/main/results/supp_data/all_raw_data.csv.

**Figure S6.**
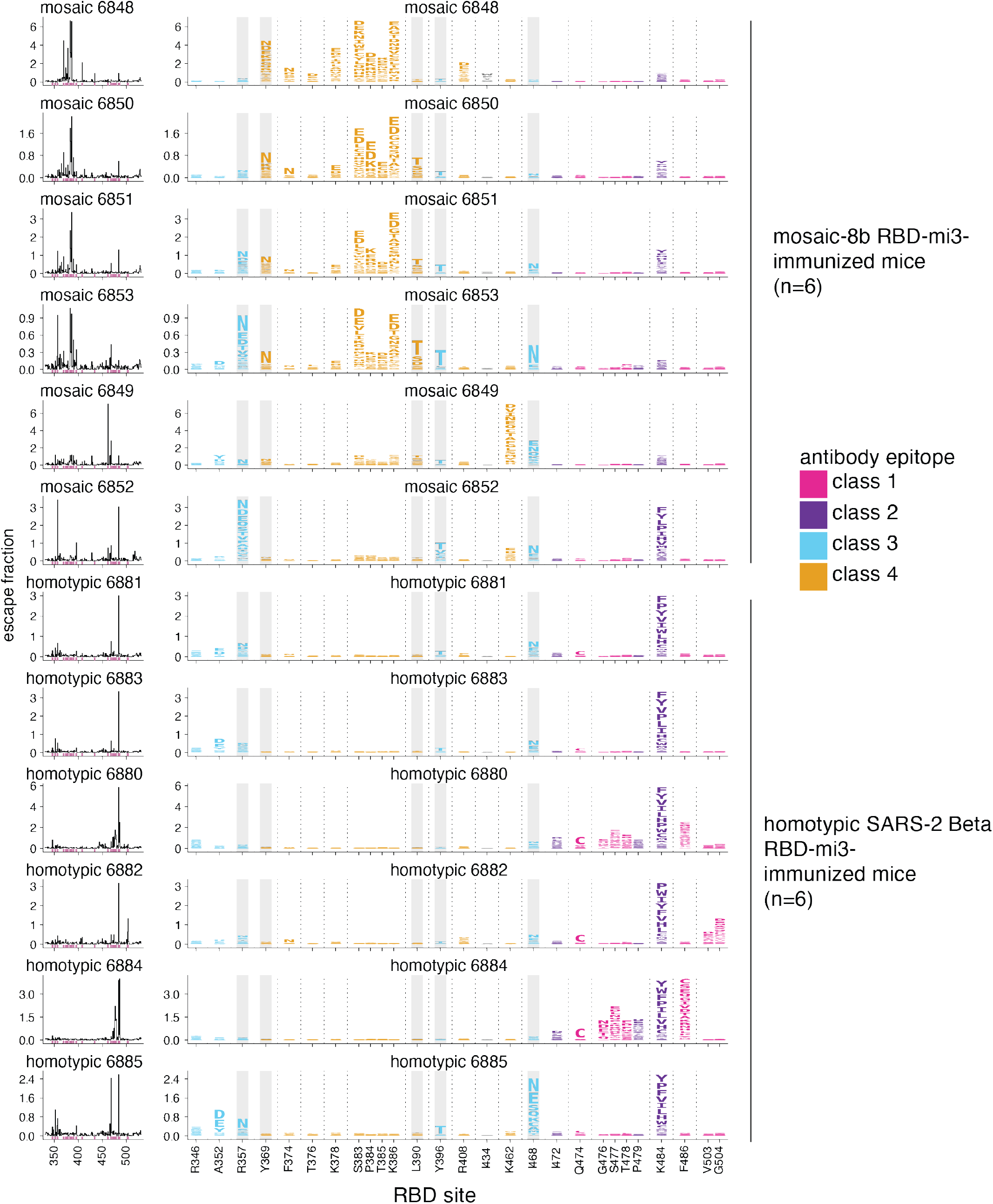
Complete antibody-escape maps for sera from mice immunized with the mosaic 8b-RBD-mi3 (top 6) or homotypic SARS-2 Beta RBD-mi3 (bottom 6) nanoparticles. The line plots at left indicate the sum of effects of all mutations at each RBD site on antibody binding, with larger values indicating more escape. The logo plots at right show key sites where mutations disrupted antibody binding (highlighted in purple on the line plot x-axes). The height of each letter is that mutation’s escape fraction. The y-axis is scaled independently for each sample. RBD sites are colored by antibody epitope, indicated at right. Sites where some mutations introduce a potential N-linked glycosylation site sequon (NxS/T) are highlighted in gray. All escape scores are in Data S2 and at https://github.com/jbloomlab/SARS-CoV-2-RBD_Beta_mosaic_np_vaccine/blob/main/results/supp_data/all_raw_data.csv. Interactive versions of logo plots and structural visualizations are at https://jbloomlab.github.io/SARS-CoV-2-RBD_Beta_mosaic_np_vaccine/.

**Figure S7.**
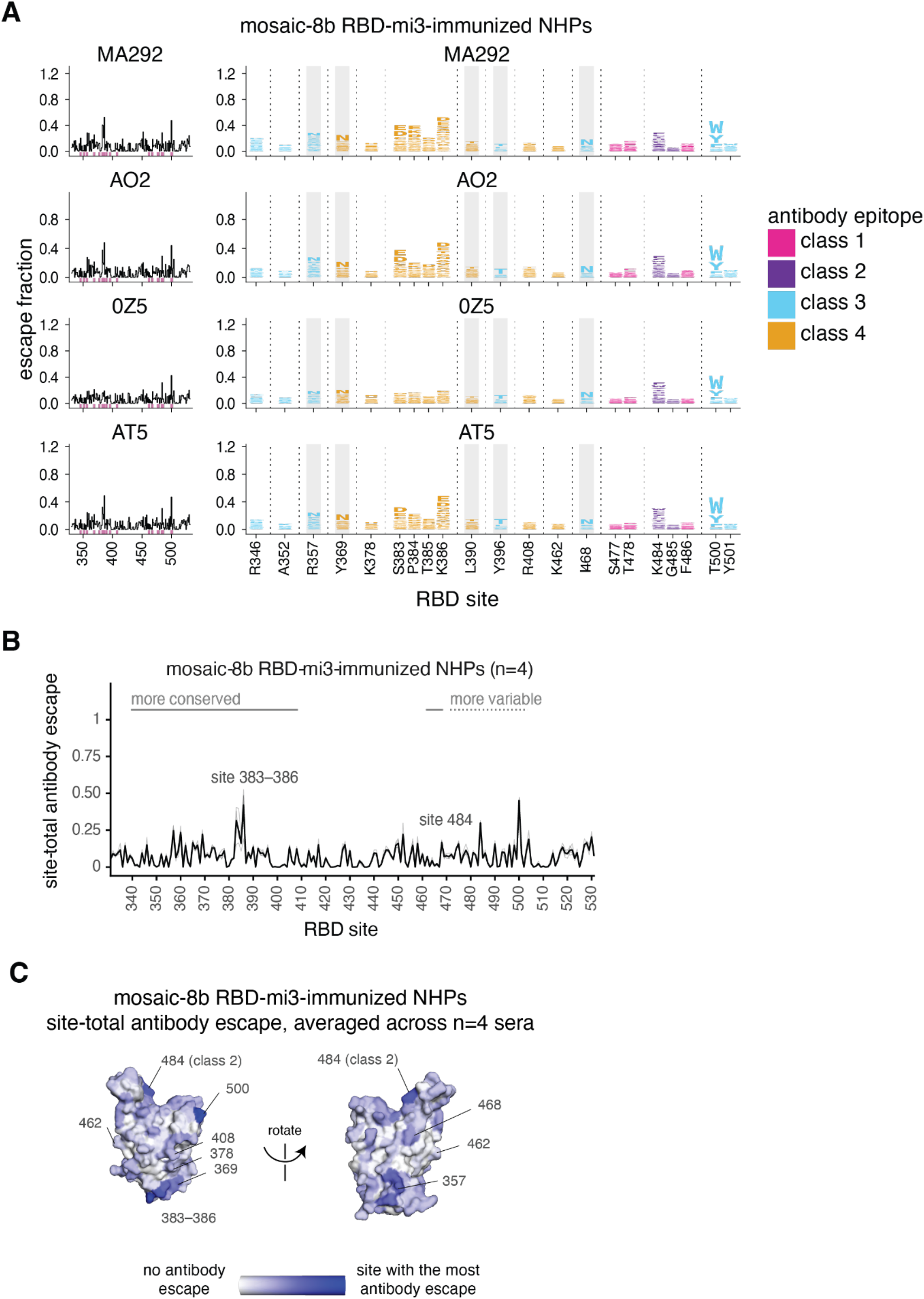
Complete antibody-escape maps for sera from NHPs immunized with mosaic 8b-RBD-mi3. (**A**) As in fig. S6, line plots (left) and logo plots (right) indicate the sum of the escape fractions for each mutation at a site, or mutation-level escape fractions for key sites, respectively. The y-axis is scaled independently for each sample. Sites where mutations introduce a potential N-linked glycosylation site sequon (NxS/T) are highlighted in gray. RBD sites are colored by antibody epitope, indicated in panel B. **(B)** The site-total antibody escape is averaged across n=4 sera, with the y-axis scaled as in panel A. (C) The average site-total antibody escape is mapped to the surface of the SARS-2 Beta RBD (PDB 7LYQ), with white indicating no escape, and blue indicating the site with the most antibody escape. Key sites are labeled, and labels are colored by antibody class. All escape scores are in Data S2 and at https://github.com/jbloomlab/SARS-CoV-2-RBD_Beta_mosaic_np_vaccine/blob/main/results/supp_data/all_raw_data.csv. Interactive versions of logo plots and structural visualizations are at https://jbloomlab.github.io/SARS-CoV-2-RBD_Beta_mosaic_np_vaccine/.

**Table S1.** Pathology and immunohistochemistry (IHC) for lung tissue isolated from vaccinated K18-hACE2 mice challenged with either SARS-2 Beta or SARS-1. Scoring for hematoxylin and eosin (H&E) is as follows: 0 = not present; 1 = minimal, 1-10%; 2 = mild, 11-25%; 3 = moderate, 26-50%; 4 = marked, 51-75%; 5 = severe, 76-100%. Scoring for IHC is as follows: 0 = not present; 1 = rare/few; 2 = scattered; 3 = moderate; 4 = numerous; 5 = diffuse. Each column represents a single animal.

|  |  | SARS-2 Beta |  |  |  |  |  |  |  |  |  |  |  |  |  |  |  |
| --- | --- | --- | --- | --- | --- | --- | --- | --- | --- | --- | --- | --- | --- | --- | --- | --- | --- |
| H & E Staining | Lung | Mosaic-8b |  |  |  | Mosaic-8gm |  |  |  | Homotypic SARS-2 Beta |  |  |  | mi3 control |  |  |  |
|  | Lesions % | 0 | 1 | 0 | 0 | 0 | 0 | 5 | 20 | 0 | 0 | 0 | 0 | <1 | 30 | 5 | 0 |
|  | Interstitial pneumonia | 0 | 0 | 0 | 0 | 0 | 0 | 1 | 2 | 0 | 0 | 0 | 0 | 0 | 1 | 1 | 0 |
|  | Bronchiolitis | 0 | 1 | 0 | 0 | 0 | 0 | 1 | 2 | 0 | 0 | 0 | 0 | 0 | 1 | 1 | 0 |
|  | perivascular leukocyte cuffing | 0 | 0 | 0 | 0 | 0 | 0 | 1 | 2 | 0 | 0 | 0 | 1 | 1 | 3 | 1 | 0 |
| IHC | Staining % | 0 | 5 | 0 | <1 | 0 | <1 | 10 | 40 | 0 | <1 | <1 | 0 | <1 | 90 | 20 | 0 |
|  | Type I and II pneumocytes | 0 | 2 | 0 | 2 | 0 | 1 | 3 | 4 | 0 | 1 | 1 | 0 | 1 | 5 | 2 | 0 |

**Table S1.** Pathology and immunohistochemistry (IHC) for lung tissue isolated from vaccinated K18-hACE2 mice challenged with either SARS-2 Beta or SARS-1.
|  |  | SARS-1 |  |  |  |  |  |  |  |  |  |  |  |  |  |  |  |
| --- | --- | --- | --- | --- | --- | --- | --- | --- | --- | --- | --- | --- | --- | --- | --- | --- | --- |
| H & E Staining | Lung | Mosaic-8b |  |  |  | Mosaic-8gm |  |  |  | Homotypic SARS-2 Beta |  |  |  | mi3 control |  |  |  |
|  | Lesions % | 0 | 0 | 0 | 0 | 0 | 0 | <1 | 0 | 0 | 0 | 0 | 0 | 0 | 5 | 0 | 0 |
|  | Interstitial pneumonia | 0 | 0 | 0 | 0 | 0 | 0 | 1 | 0 | 0 | 0 | 0 | 0 | 0 | 1 | 0 | 0 |
|  | Bronchiolitis | 0 | 0 | 0 | 0 | 0 | 0 | 1 | 0 | 0 | 0 | 0 | 0 | 0 | 1 | 0 | 0 |
|  | perivascular leukocyte cuffing | 0 | 0 | 0 | 0 | 0 | 0 | 0 | 1 | 0 | 0 | 1 | 0 | 0 | 0 | 1 | 0 |
| IHC | Staining % | <1 | <1 | 0 | 0 | 0 | <1 | 0 | 0 | 0 | 0 | <1 | 0 | 10 | 90 | 60 | 50 |
|  | Type I and II pneumocytes | 2 | 1 | 0 | 0 | 0 | 2 | 0 | 0 | 0 | 0 | 2 | 0 | 2 | 5 | 4 | 4 |

## References

1. C. Zheng et al., Real-world effectiveness of COVID-19 vaccines: a literature review and meta-analysis. Int J Infect Dis 114, 252–260 (2022).

2. D. Planas et al., Sensitivity of infectious SARS-CoV-2 B.1.1.7 and B.1.351 variants to neutralizing antibodies. Nat Med 27, 917–924 (2021).

3. N. L. Washington et al., Emergence and rapid transmission of SARS-CoV-2 B.1.1.7 in the United States. Cell 184, 2587–2594 e2587 (2021).

4. T. K. Burki, Omicron variant and booster COVID-19 vaccines. The Lancet Respiratory Medicine, (2021).

5. L. Liu et al., Striking Antibody Evasion Manifested by the Omicron Variant of SARS-CoV-2. Nature, (2021).

6. D. Yamasoba et al., Virological characteristics of SARS-CoV-2 BA.2 variant. bioRxiv, (2022).

7. F. Konings et al., SARS-CoV-2 Variants of Interest and Concern naming scheme conducive for global discourse. Nat Microbiol 6, 821–823 (2021).

8. S. Alkhovsky et al., SARS-like Coronaviruses in Horseshoe Bats (Rhinolophus spp.) in Russia, 2020. Viruses 14, (2022).

9. D. Delaune et al., A novel SARS-CoV-2 related coronavirus in bats from Cambodia. Nat Commun 12, 6563 (2021).

10. H. Zhou et al., Identification of novel bat coronaviruses sheds light on the evolutionary origins of SARS-CoV-2 and related viruses. bioRxiv, (2021).

11. S. Wacharapluesadee et al., Evidence for SARS-CoV-2 related coronaviruses circulating in bats and pangolins in Southeast Asia. Nat Commun 12, 972 (2021).

12. S. N. Seifert, M. C. Letko, A sarbecovirus found in Russian bats uses human ACE2. bioRxiv, (2021).

13. T. N. Starr et al., ACE2 binding is an ancestral and evolvable trait of sarbecoviruses. Nature, (2022).

14. M. Letko, A. Marzi, V. Munster, Functional assessment of cell entry and receptor usage for SARS-CoV-2 and other lineage B betacoronaviruses. Nature Microbiology 5, 562–569 (2020).

15. T. S. Fung, D. X. Liu, Human Coronavirus: Host-Pathogen Interaction. Annu Rev Microbiol 73, 529–557 (2019).

16. P. J. M. Brouwer et al., Potent neutralizing antibodies from COVID-19 patients define multiple targets of vulnerability. Science 369, 643–650 (2020).

17. Y. Cao et al., Potent neutralizing antibodies against SARS-CoV-2 identified by high-throughput single-cell sequencing of convalescent patients’ B cells. Cell, (2020).

18. C. Kreer et al., Longitudinal Isolation of Potent Near-Germline SARS-CoV-2-Neutralizing Antibodies from COVID-19 Patients. Cell, (2020).

19. L. Liu et al., Potent neutralizing antibodies against multiple epitopes on SARS-CoV-2 spike. Nature 584, 450–456 (2020).

20. D. F. Robbiani et al., Convergent antibody responses to SARS-CoV-2 in convalescent individuals. Nature 584, 437–442 (2020).

21. R. Shi et al., A human neutralizing antibody targets the receptor-binding site of SARS-CoV-2. Nature 584, 120–124 (2020).

22. S. J. Zost et al., Rapid isolation and profiling of a diverse panel of human monoclonal antibodies targeting the SARS-CoV-2 spike protein. Nat Med 26, 1422–1427 (2020).

23. T. F. Rogers et al., Isolation of potent SARS-CoV-2 neutralizing antibodies and protection from disease in a small animal model. Science 369, 956–963 (2020).

24. E. Seydoux et al., Analysis of a SARS-CoV-2-Infected Individual Reveals Development of Potent Neutralizing Antibodies with Limited Somatic Mutation. Immunity 53, 98–105 e105 (2020).

25. S. J. Zost et al., Potently neutralizing and protective human antibodies against SARS-CoV-2. Nature 584, 443–449 (2020).

26. C. O. Barnes et al., SARS-CoV-2 neutralizing antibody structures inform therapeutic strategies. Nature 588, 682–687 (2020).

27. D. Pinto et al., Cross-neutralization of SARS-CoV-2 by a human monoclonal SARS-CoV antibody. Nature 583, 290–295 (2020).

28. L. Piccoli et al., Mapping neutralizing and immunodominant sites on the SARS-CoV-2 spike receptor-binding domain by structure-guided high-resolution serology. Cell, (2020).

29. Z. Wang et al., mRNA vaccine-elicited antibodies to SARS-CoV-2 and circulating variants. Nature 592, 616–622 (2021).

30. H. Kleanthous et al., Scientific rationale for developing potent RBD-based vaccines targeting COVID-19. NPJ Vaccines 6, 128 (2021).

31. H. Liu et al., Cross-Neutralization of a SARS-CoV-2 Antibody to a Functionally Conserved Site Is Mediated by Avidity. Immunity 53, 1272–1280.e1275 (2020).

32. C. A. Jette et al., Broad cross-reactivity across sarbecoviruses exhibited by a subset of COVID-19 donor-derived neutralizing antibodies. Cell reports, in press (2021).

33. D. L. Burnett et al., Immunizations with diverse sarbecovirus receptor-binding domains elicit SARS-CoV-2 neutralizing antibodies against a conserved site of vulnerability. Immunity 54, 2908–2921 e2906 (2021).

34. A. A. Cohen et al., Mosaic nanoparticles elicit cross-reactive immune responses to zoonotic coronaviruses in mice. Science 371, 735–741 (2021).

35. T. K. Tan et al., A COVID-19 vaccine candidate using SpyCatcher multimerization of the SARS-CoV-2 spike protein receptor-binding domain induces potent neutralising antibody responses. Nat Commun 12, 542 (2021).

36. K. D. Brune et al., Plug-and-Display: decoration of Virus-Like Particles via isopeptide bonds for modular immunization. Scientific reports 6, 19234 (2016).

37. B. Zakeri et al., Peptide tag forming a rapid covalent bond to a protein, through engineering a bacterial adhesin. Proc Natl Acad Sci U S A 109, E690–697 (2012).

38. A. H. Keeble et al., Approaching infinite affinity through engineering of peptide–protein interaction. Proceedings of the National Academy of Sciences 116, 26523–26533 (2019).

39. A. M. Davidson, J. Wysocki, D. Batlle, Interaction of SARS-CoV-2 and Other Coronavirus With ACE (Angiotensin-Converting Enzyme)-2 as Their Main Receptor: Therapeutic Implications. Hypertension 76, 1339–1349 (2020).

40. W. Dong et al., The K18-Human ACE2 Transgenic Mouse Model Recapitulates Non-severe and Severe COVID-19 in Response to an Infectious Dose of the SARS-CoV-2 Virus. J Virol 96, e0096421 (2022).

41. C. K. Yinda et al., K18-hACE2 mice develop respiratory disease resembling severe COVID-19. PLoS Pathog 17, e1009195 (2021).

42. E. S. Winkler et al., SARS-CoV-2 infection of human ACE2-transgenic mice causes severe lung inflammation and impaired function. Nat Immunol 21, 1327–1335 (2020).

43. C. L. Hsieh et al., Structure-based design of prefusion-stabilized SARS-CoV-2 spikes. Science 369, 1501–1505 (2020).

44. A. J. Greaney et al., Comprehensive mapping of mutations in the SARS-CoV-2 receptor-binding domain that affect recognition by polyclonal human plasma antibodies. Cell Host Microbe 29, 463–476 e466 (2021).

45. T. Zohar et al., Compromised Humoral Functional Evolution Tracks with SARS-CoV-2 Mortality. Cell 183, 1508–1519 e1512 (2020).

46. J. Yu et al., DNA vaccine protection against SARS-CoV-2 in rhesus macaques. Science 369, 806–811 (2020).

47. N. van Doremalen et al., Intranasal ChAdOx1 nCoV-19/AZD1222 vaccination reduces viral shedding after SARS-CoV-2 D614G challenge in preclinical models. Science translational medicine 13, (2021).

48. G. Dagotto et al., Comparison of Subgenomic and Total RNA in SARS-CoV-2 Challenged Rhesus Macaques. J Virol, (2021).

49. P. Kumari et al., Neuroinvasion and Encephalitis Following Intranasal Inoculation of SARS-CoV-2 in K18-hACE2 Mice. Viruses 13, (2021).

50. B. Pulendran, S. A. P, D. T. O’Hagan, Emerging concepts in the science of vaccine adjuvants. Nat Rev Drug Discov 20, 454–475 (2021).

51. P. B. Gilbert et al., Immune correlates analysis of the mRNA-1273 COVID-19 vaccine efficacy clinical trial. Science, eab3435 (2021).

52. F. Hansen et al., SARS-CoV-2 reinfection prevents acute respiratory disease in Syrian hamsters but not replication in the upper respiratory tract. Cell reports, (2022).

53. A. J. Greaney et al., Complete Mapping of Mutations to the SARS-CoV-2 Spike Receptor-Binding Domain that Escape Antibody Recognition. Cell Host Microbe 29, 44–57 e49 (2021).

54. A. J. Greaney et al., A SARS-CoV-2 variant elicits an antibody response with a shifted immunodominance hierarchy. PLoS Pathog 18, e1010248 (2022).

55. A. C. Walls et al., Distinct sensitivities to SARS-CoV-2 variants in vaccinated humans and mice. bioRxiv, (2022).

56. R. Shinnakasu et al., Glycan engineering of the SARS-CoV-2 receptor-binding domain elicits cross-neutralizing antibodies for SARS-related viruses. J Exp Med 218, (2021).

57. J. Heeney et al., Gene delivery of a single, structurally engineered Coronavirus vaccine antigen elicits SARS-CoV-2 Omicron and pan-Sarbecovirus neutralisation. Research Square, (2021).

58. W. Dejnirattisai et al., The antigenic anatomy of SARS-CoV-2 receptor binding domain. Cell, (2021).

59. A. J. Greaney et al., Mapping mutations to the SARS-CoV-2 RBD that escape binding by different classes of antibodies. Nat Commun 12, 4196 (2021).

60. T. N. Starr et al., SARS-CoV-2 RBD antibodies that maximize breadth and resistance to escape. Nature 597, 97–102 (2021).

61. T. N. Starr et al., Prospective mapping of viral mutations that escape antibodies used to treat COVID-19. Science 371, 850–854 (2021).

62. E. Cameroni et al., Broadly neutralizing antibodies overcome SARS-CoV-2 Omicron antigenic shift. Nature, (2021).

63. D. J. Sheward et al., Structural basis of Omicron neutralization by affinity-matured public antibodies. bioRxiv, (2022).

64. K. Westendorf et al., LY-CoV1404 (bebtelovimab) potently neutralizes SARS-CoV-2 variants. bioRxiv, (2022).

65. M. G. Joyce et al., SARS-CoV-2 ferritin nanoparticle vaccines elicit broad SARS coronavirus immunogenicity. Cell reports 37, 110143 (2021).

66. K. O. Saunders et al., Neutralizing antibody vaccine for pandemic and pre-emergent coronaviruses. Nature 594, 553–559 (2021).

67. A. E. Powell et al., A Single Immunization with Spike-Functionalized Ferritin Vaccines Elicits Neutralizing Antibody Responses against SARS-CoV-2 in Mice. ACS Cent Sci 7, 183–199 (2021).

68. P. T. Heath et al., Safety and Efficacy of NVX-CoV2373 Covid-19 Vaccine. N Engl J Med 385, 1172–1183 (2021).

69. W. Wang, B. Huang, Y. Zhu, W. Tan, M. Zhu, Ferritin nanoparticle-based SARS-CoV-2 RBD vaccine induces a persistent antibody response and long-term memory in mice. Cell Mol Immunol 18, 749–751 (2021).

70. X. Ma et al., Nanoparticle Vaccines Based on the Receptor Binding Domain (RBD) and Heptad Repeat (HR) of SARS-CoV-2 Elicit Robust Protective Immune Responses. Immunity 53, 1315–1330.e1319 (2020).

71. Q. Geng et al., Novel virus-like nanoparticle vaccine effectively protects animal model from SARS-CoV-2 infection. PLoS Pathog 17, e1009897 (2021).

72. Y. F. Kang et al., Rapid Development of SARS-CoV-2 Spike Protein Receptor-Binding Domain Self-Assembled Nanoparticle Vaccine Candidates. ACS Nano 15, 2738–2752 (2021).

73. A. C. Walls et al., Elicitation of Potent Neutralizing Antibody Responses by Designed Protein Nanoparticle Vaccines for SARS-CoV-2. Cell 183, 1367–1382 e1317 (2020).

74. D. Li et al., Breadth of SARS-CoV-2 Neutralization and Protection Induced by a Nanoparticle Vaccine. bioRxiv, (2022).

75. A. I. Mosa, Antigenic Variability. Front Immunol 11, 2057 (2020).

76. M. Kanekiyo et al., Mosaic nanoparticle display of diverse influenza virus hemagglutinins elicits broad B cell responses. Nat Immunol 20, 362–372 (2019).

77. A. C. Walls et al., Elicitation of broadly protective sarbecovirus immunity by receptor-binding domain nanoparticle vaccines. Cell, (2021).

78. T. U. J. Bruun, A. C. Andersson, S. J. Draper, M. Howarth, Engineering a Rugged Nanoscaffold To Enhance Plug-and-Display Vaccination. ACS Nano 12, 8855–8866 (2018).

79. M. Landau et al., ConSurf 2005: the projection of evolutionary conservation scores of residues on protein structures. Nucleic Acids Res 33, W299–302 (2005).

80. S. Guindon et al., New algorithms and methods to estimate maximum-likelihood phylogenies: assessing the performance of PhyML 3.0. Syst Biol 59, 307–321 (2010).

81. F. Sievers et al., Fast, scalable generation of high-quality protein multiple sequence alignments using Clustal Omega. Molecular Systems Biology 7, 539 (2011).

82. C. O. Barnes et al., Structures of Human Antibodies Bound to SARS-CoV-2 Spike Reveal Common Epitopes and Recurrent Features of Antibodies. Cell 182, 828–842 e816 (2020).

83. C.-L. Hsieh et al., Structure-based Design of Prefusion-stabilized SARS-CoV-2 Spikes. bioRxiv, (2020).

84. A. A. Cohen et al., Construction, characterization, and immunization of nanoparticles that display a diverse array of influenza HA trimers. PLoS One 16, e0247963 (2021).

85. L. J. Reed, H. Muench, A Simple Method of Estimating Fifty Per Cent Endpoints12. American Journal of Epidemiology 27, 493–497 (1938).

86. K. H. D. Crawford et al., Protocol and Reagents for Pseudotyping Lentiviral Particles with SARS-CoV-2 Spike Protein for Neutralization Assays. Viruses 12, 513 (2020).

87. A. P. West, Jr. et al., Computational analysis of anti-HIV-1 antibody neutralization panel data to identify potential functional epitope residues. Proc Natl Acad Sci U S A 110, 10598–10603 (2013).

88. T. N. Starr et al., Deep Mutational Scanning of SARS-CoV-2 Receptor Binding Domain Reveals Constraints on Folding and ACE2 Binding. Cell 182, 1295–1310.e1220 (2020).

89. S. K. Hilton, et al., dms-view: Interactive visualization tool for deep mutational scanning data. J Open Source Softw 5, (2020).

