## Supplementary material for "Mosaic RBD nanoparticles protect against multiple sarbecovirus challenges in animal models": Pathology summary

**I. Study MS21-171 (ACUC2021-032): Efficacy of Mosaic-8 nanoparticle vaccine against SARS-CoV-2 or SARS-CoV in mice**

**II. Data**

The data set includes an Excel spreadsheet with lesion severity scores for each animal and figure submission for a manuscript.

**III. Materials and Methods**

Tissues were fixed in 10% Neutral Buffered Formalin x2 changes, for a minimum of 7 days. Tissues were placed in cassettes and processed with a Sakura VIP-6 Tissue Tek, on a 12-hour automated schedule, using a graded series of ethanol, xylene, and PureAffin. Embedded tissues are sectioned at 5um and dried overnight at 42 degrees C prior to staining. Specific anti-CoV immunoreactivity was detected using SARS-CoV-2 nucleocapsid antibody (Genscript) at a 1:1000 dilution. The secondary antibody is the Vector Laboratories ImPress VR anti-rabbit IgG polymer (cat# MP-6401). The tissues were then processed for immunohistochemistry using the Discovery Ultra automated stainer (Roche Tissue Diagnostics) with a ChromoMap DAB kit (Roche Tissue Diagnostics cat#760-159).

**IV. Summary**

This study is composed of 32 K18-hACE2 mice.

Study is designed to investigate whether the vaccine is able to protect mice against SARS-CoV-2 infection, which is found on the nanoparticle, as well as against SARS-CoV infection, which is not found on the nanoparticle. As a secondary objective, we will include a group which will be vaccinated with Mosaic-8-N, which contains an N glycan at amino acid 484 in the spike protein. This amino acid is often substituted in variants of SARS-CoV-2 that are able to evade the immune system, and we hypothesize that including an N-glycan site at this position will broaden the immune response. Comparisons will be done against an unconjugated nanoparticle vaccine and a SARS-CoV-2 homotypic nanoparticle vaccine.

Hypothesis: Mosaic-8 vaccine protects mice against a lethal infection with SARS-CoV-2 or SARS-CoV

Aim 1: To determine the efficacy of Mosaic-8 and Mosaic-8-N against SARS-CoV-2

Aim 2: To determine the efficacy of Mosaic-8 and Mosaic-8-N against SARS-CoV

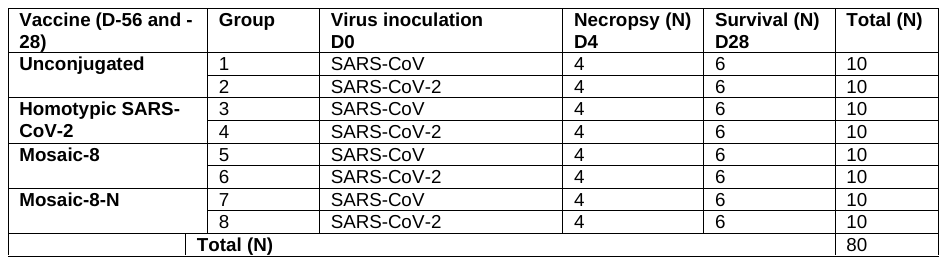

**Histopathology**

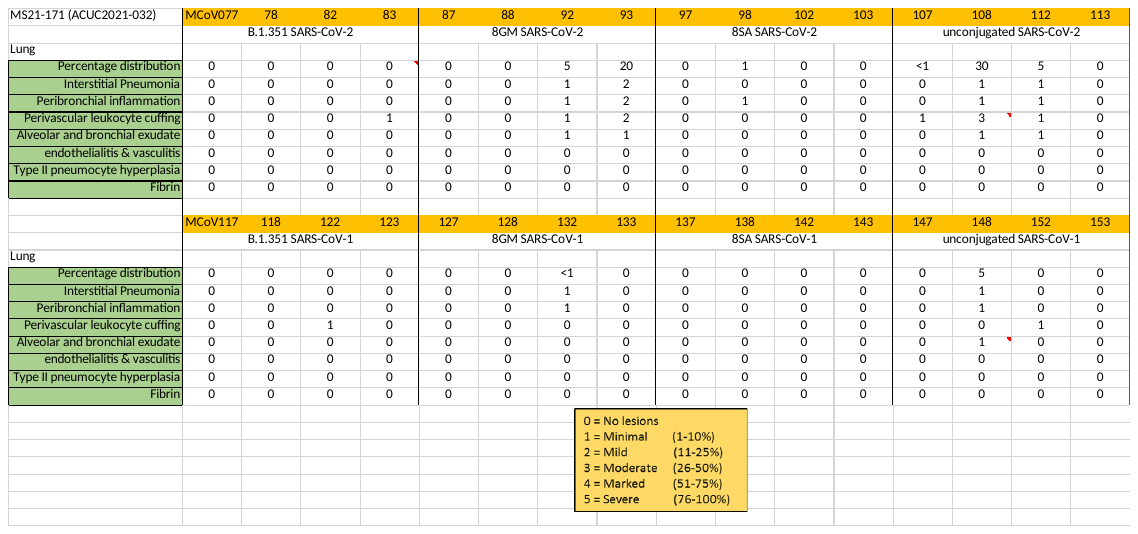

H&E Control animals

Control animals were necropsied on D4. SARS-CoV-2 lesions were evident in two of four animals affecting between 5-30% of the lung. Lesions are characterized by minimal interstitial pneumonia centered on terminal bronchioles and extending into the adjacent alveoli with minimal-moderate perivascular leucocyte cuffing and alveolar exudate. One animal in the SARS-CoV control group had a lesion affecting 5% of the lung and is characterized by a minimal interstitial pneumonia centered on terminal bronchioles with minimal alveolar exudate.

H&E Vaccinated animals exposed to SARS-CoV-2

Animals were vaccinated with Mosaic-8b, Mosaic-8gm or Homotypic SARS-2 Beta and exposed to SARS-CoV-2. Mosaic-8b vaccinated animals did not have interstitial pneumonia and only one animal had minimal peribronchial inflammation. Two of four animals vaccinated with Moisac-8gm had minimal to mild interstitial pneumonia affecting between 5-20% of the lungs. Animals vaccinated with Homotypic SARS-2 Beta vaccine did not have pulmonary lesions except for one animal with minimal perivascular cuffing.

H&E Vaccinated animals exposed to SARS-CoV

Animals were vaccinated with Mosaic-8b, Mosaic-8gm or Homotypic SARS-2 Beta and exposed to SARS-CoV-2. Mosaic-8b vaccinated animals did not have pulmonary pathology. One animal vaccinated with Moisac-8gm had minimal interstitial pneumonia with peribronchial inflammation affecting less than 1% of the lung. Animals vaccinated with Homotypic SARS-2 Beta vaccine did not have pulmonary lesions except for one animal with minimal perivascular cuffing.

**Immunohistochemistry**

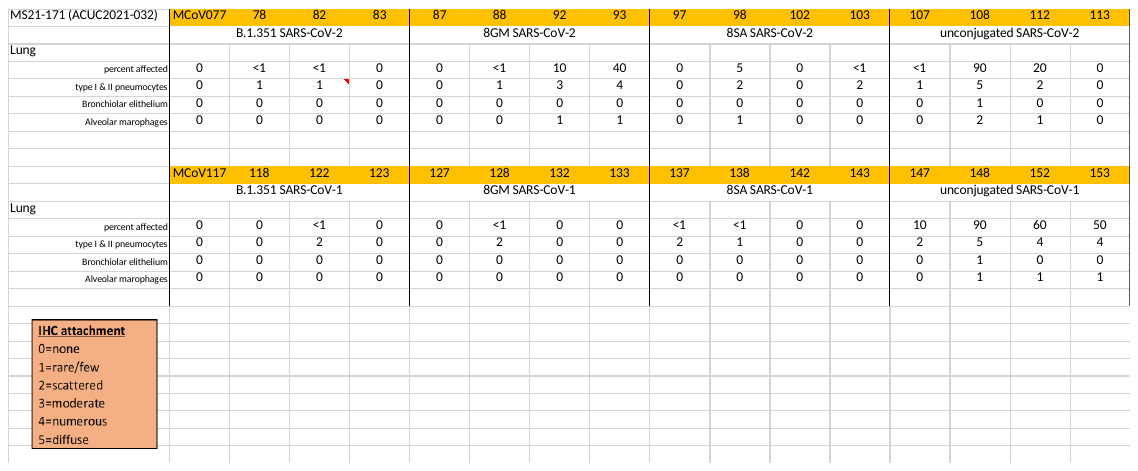

IHC Control animals

Control animals were necropsied on D4. SARS-CoV-2 immunoreactivity is evident in three of four animals affecting between 1-90% of the lung. Antigen is observed predominately in type I and II pneumocytes as well as fewer pulmonary macrophages and rarely in bronchiolar epithelium in one animal. The SARS-CoV control group expressed immunoreactivity in in all animals affecting between 10-90% of the lung. Antigen is observed predominately in type I and II pneumocytes as well as fewer pulmonary macrophages and rarely in bronchiolar epithelium in one animal.

IHC Vaccinated animals exposed to SARS-CoV2

Two of four animals vaccinated with Mosaic-8b expressed antigen in Type 1&2 pneumocytes in 1-5% of the lung. Vaccination with Mosaic-8gm also resulted in antigen expression in three of four animals in 1-40% of the lung. Vaccination with Homotypic SARS-2 Beta resulted in two of four animals with less than 1% of the lung affected.

IHC Vaccinated animals exposed to SARS-CoV

Two of four animals vaccinated with Mosaic-8b expressed antigen in Type 1&2 pneumocytes in less than 1% of the lung. Vaccination with Mosaic-8gm also resulted in scattered antigen expression in one of four animals in less than 1% of the lung. Vaccination with Homotypic SARS-2 Beta resulted in one of four animals with less than 1% of the lung affected.

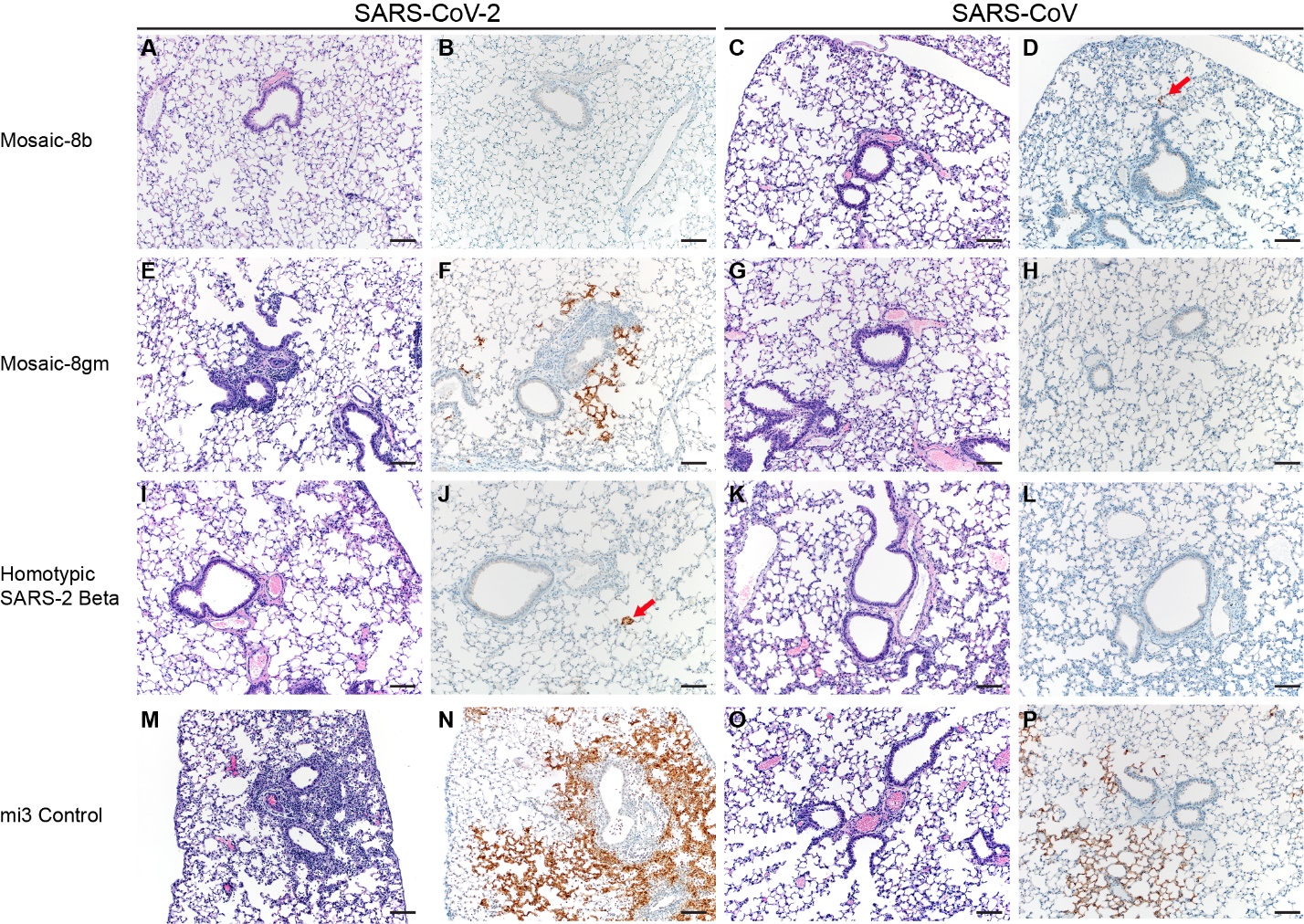

**Figure ?. Pulmonary effects of vaccinated and control K18-hACE2 mice at 4 DPI**. H&E staining (1st and 3rd column) and IHC staining against N protein (brown, 2nd and 4th column), 100x, scale bar = 100µm; E/M: minimal-mild peribronchial inflammation D/E: red arrow = immunoreactivity

Greg Saturday, DVM, DACVP, DABT
